# Apple and banana fruits produce anteiso- and iso-branched-chain esters from newly synthesized precursors

**DOI:** 10.1101/2023.10.26.564239

**Authors:** Philip Engelgau, Sumithra K. Wendakoon, Aubrey DuBois, Emily J. Mayhew, Randolph Beaudry

## Abstract

Inhibitors of acetohydroxyacid synthase, the common enzyme of branched-chain amino acid biosynthesis, were applied to ripening apple (*Malus* ×*domestica* Borkh.), banana (*Musa* spp.), and flowering quince (*Chaenomeles* ×*superba*) fruits to discern the contribution of newly synthesized precursors to branched-chain ester formation. After treatment, anteiso- and iso-branched-chain volatiles (i.e., those related to isoleucine, and valine and leucine, respectively) were observed to universally decrease in content. Fruits recovered production following exogenous feeding of branched-chain ⍺-ketoacids. Furthermore, apple and banana fruits were capable of metabolizing all three branched-chain ⍺-ketoacids to esters. Among free amino acids, only the branched-chain amino acids with correspondingly reduced branched-chain esters had a lesser concentration following treatment with inhibitor. Our results ultimately reject the hypothesis that anteiso- and iso-branched-chain esters are derived from preexisting amino acids and instead support the hypothesis that these esters are the product of *de novo* precursor biosynthesis. The novel use of these inhibitors also allowed for further investigation of branched-chain volatile biosynthesis, the citramalate synthase pathway, and the importance of precursor availability in fruits. Notably, in ‘Valery’ banana fruit, ethyl acetate and butyl acetate were found to be dependent on acetohydroxyacid synthase activity for production whereas 1-methylbutyl acetate and 1-methylbutyl butanoate (sec-branched-chain esters) were not. Inhibitor usage on apples also allowed for a sensory study that found that humans can discern the absence of 2-methylbutyl and 2-methylbutanoate esters in apple fruit. Additionally, a population genetics analysis found that there is selection pressure against apples that lack these esters.

## Introduction

For over half a century, a relationship has been known to exist between the branched-chain esters, which act as impact flavor notes for many popular fruits, and branched-chain amino acids. Specifically, 2-methylbutyl and 2-methylbutanote esters have been linked to isoleucine metabolism, 2-methylpropyl and 2-methylpropanoate esters to valine metabolism, and 3-methylbutyl and 3-methylbutanoate esters to leucine metabolism (**Fig. 1**; **Fig. 2**) (Myers, et al., 1970) (Rowan, et al., 1996) (Tressl & Drawert, 1973). However, the relationship between the three branched-chain amino acids and their respective esters has been disputed for some time.

**Figure 1.**
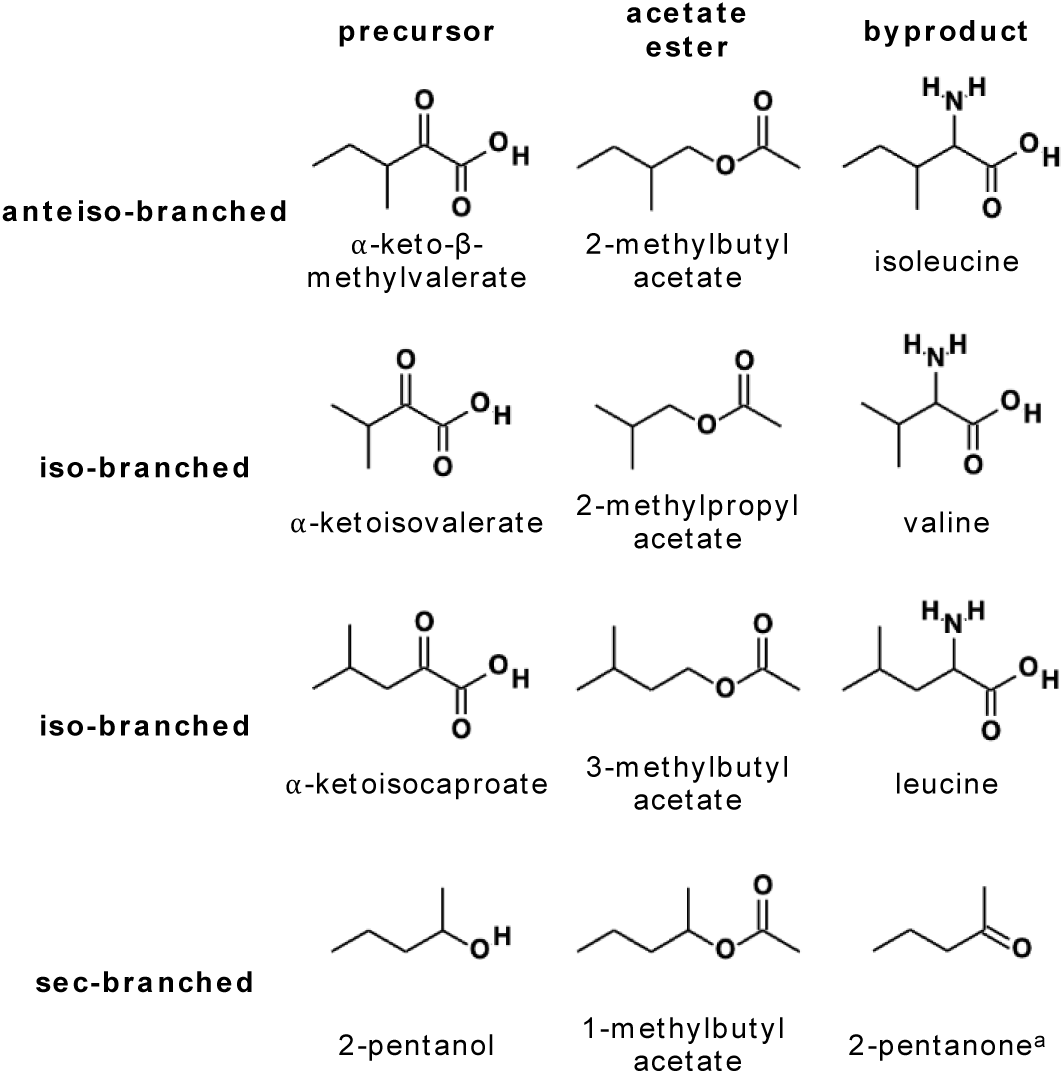
Structures of some of the precursors and byproducts of branched-chain esters discussed, including branching pattern designations. ^a^ 2-Pentanone is not technically sec-branched but is a potential oxidation byproduct of 2-pentanol.

**Figure 2.**
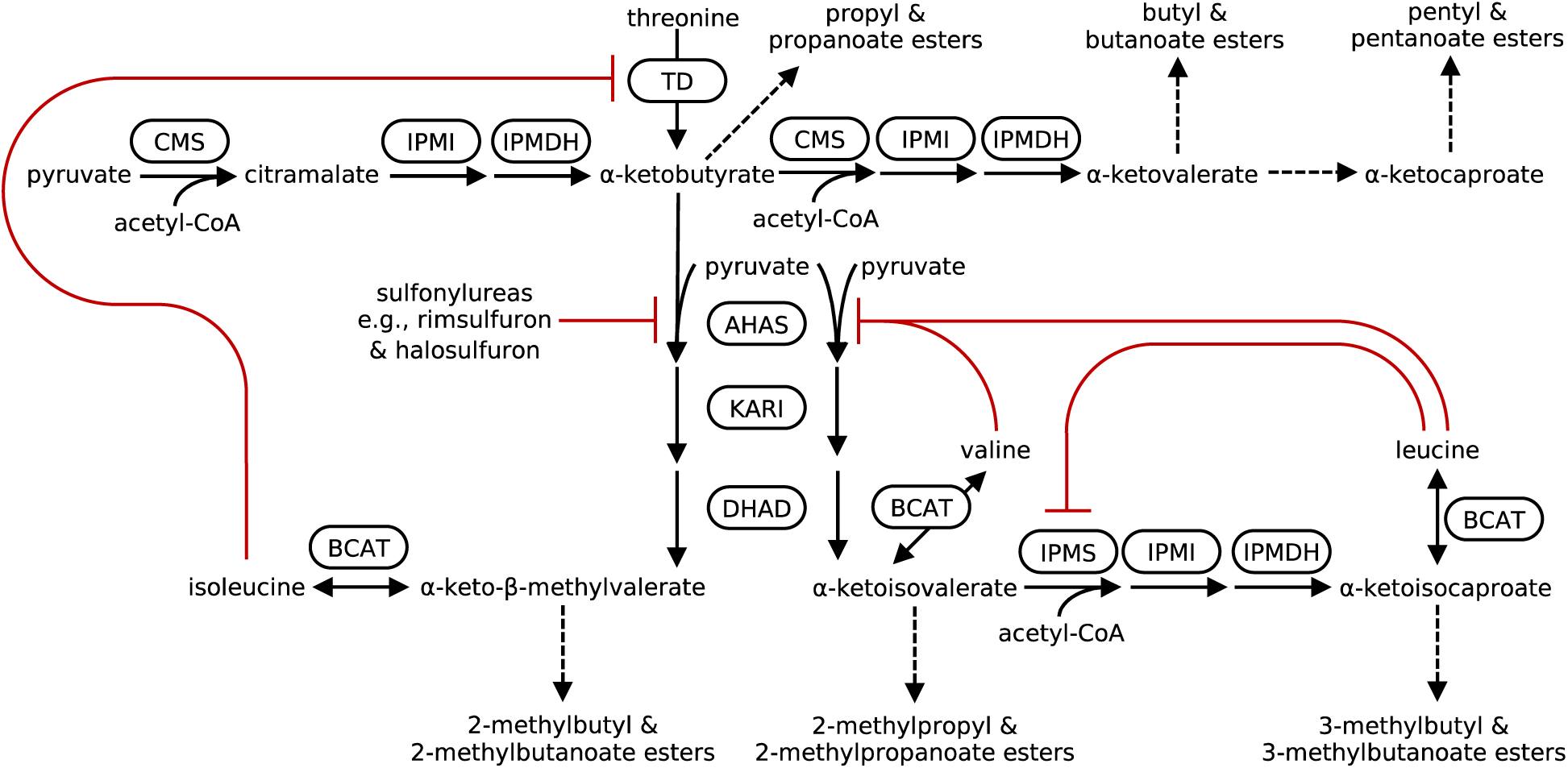
Branched-chain amino acid biosynthesis and the citramalate synthase pathway. Reactions shown as solid arrows, those understood to be freely reversible are depicted with double-ended arrows. Abbreviated pathways shown with dashed arrows. Enzymes shown in curved boxes. Principal inhibitory interactions drawn with red lines. Minor inhibitory mechanisms not shown: antagonism of isoleucine feedback at TD by valine and isoleucine inhibition of AHAS. Note that of the fruits discussed herein, CMS has only been identified in apple. Further straight-chain ⍺-ketoacid elongation via CMS has not been observed in apple fruits. For the movement of carbons as a result of labeled acetate feeding, readers are referred to Sugimoto et al., 2021. AHAS = acetohydroxyacid synthase (also known as acetolactate synthase), BCAT = branched-chain aminotransferase, CMS = citramalate synthase, DHAD = dihydroxyacid dehydratase, IPMDH = isopropylmalate dehydrogenase, IPMI = isopropylmalate isomerase, IPMS = isopropylmalate synthase, KARI = ketol-acid reductoisomerase, TD = threonine deaminase.

Based on feeding studies that have demonstrated an interchange of labeled carbons between exogenously fed branched-chain amino acids and emanated branched-chain esters, many groups have assumed that these amino acids act as direct precursors to their related volatiles via a catabolic relationship (Gonda, et al., 2010) (Rowan, et al., 1996) (Tressl & Drawert, 1973) (Myers, et al., 1970). However, such assertions overlook or disregard processes that must occur to supply said precursors. Ultimately, the idea that aroma biosynthesis results from catabolic processes, i.e., from amino acids derived via senescence-related protein degradation, has become dogma (Maoz, et al., 2022) (Kays & Paull, 2004).

Ripening in fruit is a highly dynamic process involving sequentially induced and deliberate modifications to chlorophyll content, respiration, pigmentation, starch degradation, and sugar:acid balance, to name a few (Gortner, et al., 1967). It would seem inconsistent to suggest that aroma biosynthesis, the often-terminal feature of ripening and thus the ultimate attractant for consumption and seed dispersal, is not also an active process. Our research group’s hypothesis is that aroma formation is under programmed regulation and thusly proposed that branched-chain volatiles are instead more directly derived from *de novo* synthesized precursors via active, anabolic processes.

The metabolites that directly link *de novo* branched-chain biosynthesis and branched-chain volatile production are the branched-chain α-ketoacids (**Fig. 2**). These compounds are in equilibrium with the branched-chain amino acids due to the freely reversible transamination facilitated by branched-chain amino transferase. They can also be thought of as being the last primary metabolite prior to specialization for ester synthesis. Thus, while the amino acids are more regularly measured due to the ease and commonality of said analyses, the branched-chain α-ketoacids are likely the more accurate precursor to branched-chain esters.

It has recently been observed that among the free amino acids of ripening apple (*Malus* ×*domestica* Borkh.) and banana (*Musa* spp.) fruits, only those with related branched-chain volatiles produced by the fruit undergo a marked increase that is concomitant with aroma emanation (i.e., isoleucine in apple, and valine and leucine in banana) (Alsmairat, et al., 2018) (Sugimoto, et al., 2011). Non-discriminatory protein degradative processes would not be expected to produce such coincidental results, implying that these fruits are actively engaging the synthetic processes of branched-chain amino acids, and thus branched-chain α-ketoacids.

Sugimoto et al. (2021) further demonstrated the importance of *de novo* precursor production in apple fruit through the elucidation of citramalate synthase’s role in circumventing isoleucine inhibition at threonine deaminase to produce a pool of α-keto-β-methylvalerate for 2-methylbutyl and 2-methylbutanoate ester production (**Fig. 2**). Furthermore, apple cultivars lacking a catalytically active allele of citramalate synthase were found to produce minimal quantities of said esters.

Because of the continuing controversy over the source of ester precursors, we sought to definitively determine whether branched-chain esters are synthesized from preexisting amino acids or from *de novo* biosynthesis in apple. Further, we wished to extend the inquiry beyond apple and included an ornamental flowering quince (*Chaenomeles* ×*superba*), a little explored apple relative, and banana (*Musa* spp.), a tropical monocot. We hypothesized that if ester production is dependent upon anabolic precursor synthesis in ripening fruits, then targeted inhibition of the canonical biosynthetic pathway for branched-chain α-ketoacids should simultaneously prevent the accumulation of the branched-chain α-ketoacids and their downstream metabolites including branched-chain amino acids and esters. On the other hand, if the branched-chain amino acids are catabolically derived from previously formed proteins, then branched-chain ester synthesis should persist or be minimally disrupted by inhibition of *de novo* synthesis.

To test these hypotheses, we treated ripening apple and flowering quince fruit peel, and banana fruit pulp, the sites of aroma biogenesis (Guadagni, et al., 1971) (Alsmairat, et al., 2018), with sulfonylureas. This class of compounds are one of many chemical families that are widely used as herbicides due to their ability to inhibit the common enzyme of branched-chain amino acid biosynthesis, and thus branched-chain α-ketoacid production, acetohydroxyacid synthase (AHAS, **Fig. 2**). Sulfonylureas specifically act by binding within and obstructing the substrate channel that leads to the active site of AHAS (McCourt, et al., 2006), resulting in a loss of catalytic activity. Given the lack of an alternative biosynthetic pathway, AHAS inhibition arrests isoleucine, valine, and leucine biosynthesis and ultimately translates into severe inhibition of DNA synthesis, a halt of mitosis, and eventual plant death (Shaner & Reider, 1986). Furthermore, increased protein turnover resulting in elevated total free amino acid levels, a likely effect of acute amino acid starvation, has been observed after application (Shaner & Reider, 1986) (Goldberg & St. John, 1976).

The application of AHAS inhibitors to ripening fruits should likewise halt any *de novo* production of branched-chain amino acids, α-ketoacids, and related metabolites. The resulting effects on branched-chain volatile and amino acid content should be illuminating to the aroma biochemistry of fruits.

## Results

### Apple

#### Volatiles

The application of rimsulfuron, a sulfonylurea AHAS inhibitor, led to a significant reduction of the headspace concentrations of every 2-methylbutyl and 2-methylbutanoate ester analyzed in ‘Gala’, ‘Empire’, and ‘Jonagold’ apple fruits (**Fig 3A**; **Table 1; Supplemental Fig. S1; Supplemental Fig. S2; Supplemental Table S1**). Furthermore, 2-methylpropyl acetate, which is present in quantities under 2 nmol ᐧ L^-1^ in all three cultivars, was also reduced in the headspaces of each of the cultivars when treated (**Table 2**; **Supplemental Fig. S1; Supplemental Table S1**).

**Figure 3.**
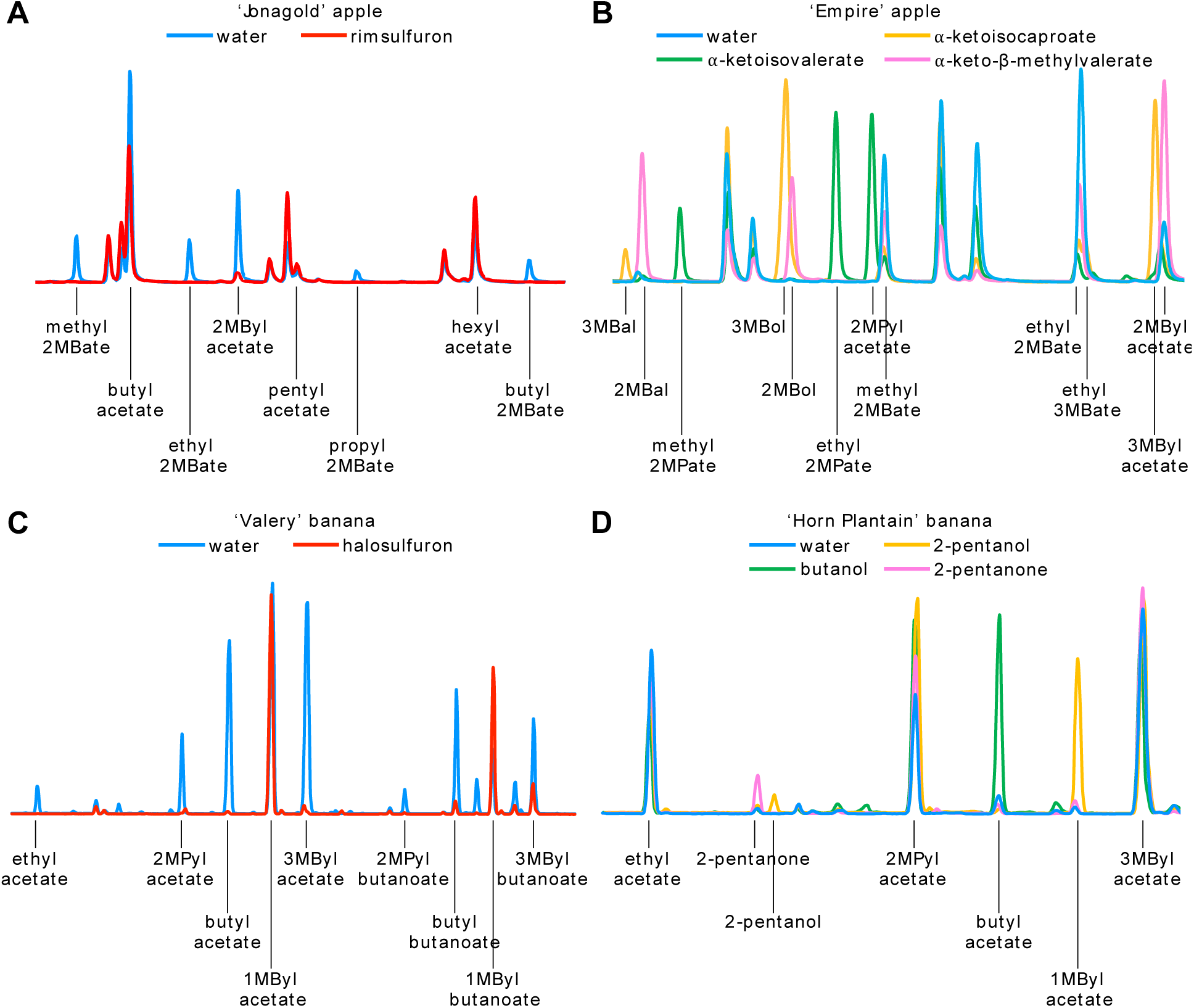
Representative sections of chromatograms from incubation chamber headspace for several of the feeding experiments discussed. A) ‘Jonagold’ apple fruit peels treated with water or rimsulfuron and fed methanol. B) ‘Empire’ apple fruit peels fed with methanol and either water or a branched-chain ⍺-ketoacid. C) ‘Valery’ banana fruit pulp sections treated with water or halosulfuron. D) ‘Horn Plantain’ banana fruit pulp sections fed with water or a potential ester precursor. 1MByl = 1-methylbutyl, 2MBal = 2-methylbutanal, 2MBate = 2-methylbutanoate, 2MBol = 2-methylbutanol, 2MByl = 2-methylbutyl, 2MPate = 2-methylpropanoate, 2MPyl = 2-methylpropyl, 3MBal = 3-methylbutanal, 3MBate = 3-methylbutanoate, 3MBol = 3-methylbutanol, 3MByl = 3-methylbutyl. For all data see Supplemental Tables and Figures.

**Table 1.**
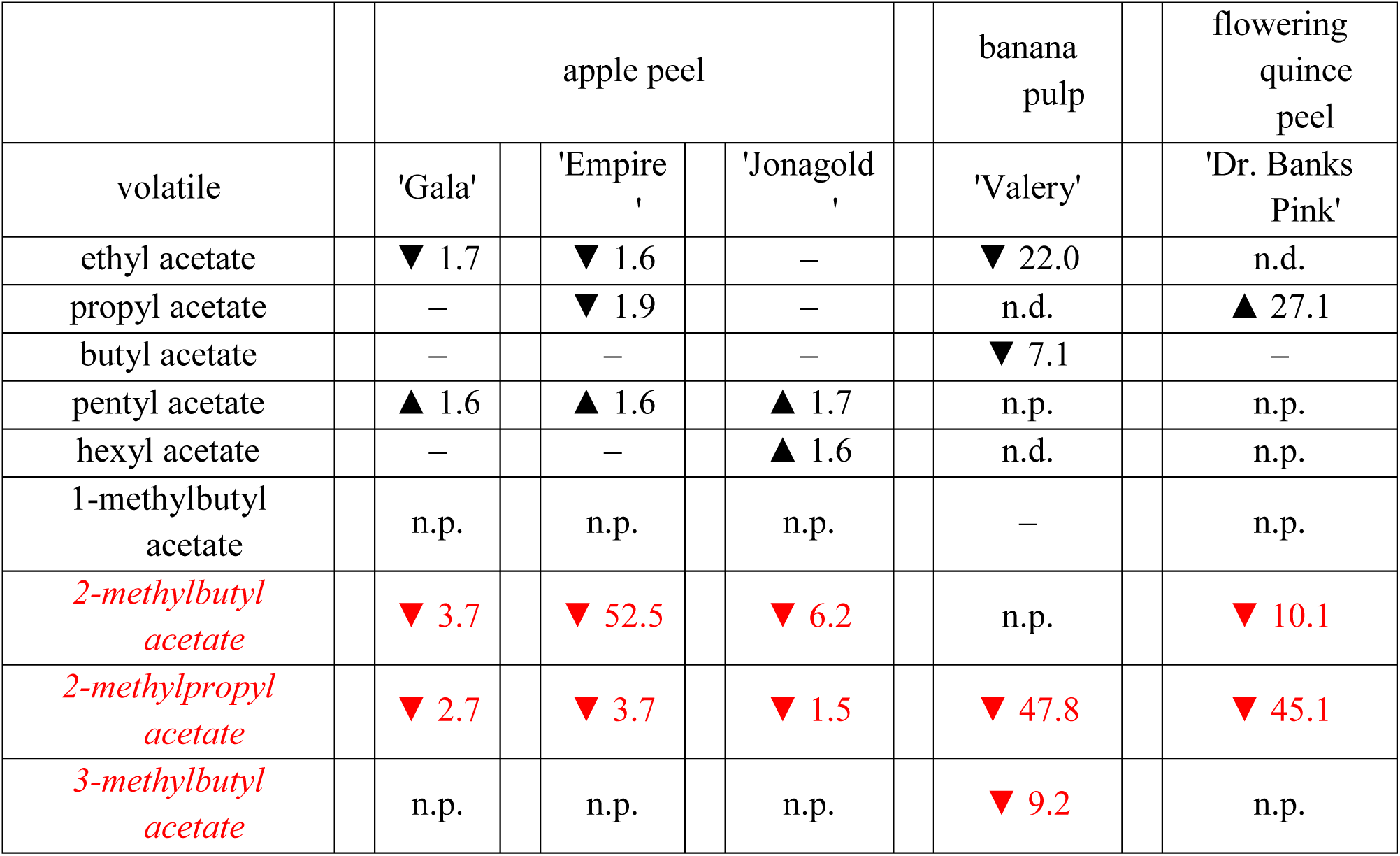
Fold-change of acetate esters of sulfonylurea-treated fruit tissues compared to control. Fold-change presented if significant (two-tailed two-sample equal variance t-test for apple and flowering quince, paired t-test for banana; α = 0.05). Anteiso- and iso-branched-chain esters highlighted in red. For quantified levels of these and other volatiles, as well as p-values, see Supplemental Figures & Tables. n.p. = not present, n.d. = not determined.

|  | apple peel |  |  | banana pulp | flowering quince peel |
| --- | --- | --- | --- | --- | --- |
| volatile | 'Gala' | 'Empire' | 'Jonagold' | 'Valery' | 'Dr. Banks Pink' |
| ethyl acetate | ▼ 1.7 | ▼ 1.6 | – | ▼ 22.0 | n.d. |
| propyl acetate | – | ▼ 1.9 | – | n.d. | ▲ 27.1 |
| butyl acetate | – | – | – | ▼ 7.1 | – |
| pentyl acetate | ▲ 1.6 | ▲ 1.6 | ▲ 1.7 | n.p. | n.p. |
| hexyl acetate | – | – | ▲ 1.6 | n.d. | n.p. |
| 1-methylbutyl acetate | n.p. | n.p. | n.p. | – | n.p. |
| *2-methylbutyl acetate* | ▼ 3.7 | ▼ 52.5 | ▼ 6.2 | n.p. | ▼ 10.1 |
| *2-methylpropyl acetate* | ▼ 2.7 | ▼ 3.7 | ▼ 1.5 | ▼ 47.8 | ▼ 45.1 |
| *3-methylbutyl acetate* | n.p. | n.p. | n.p. | ▼ 9.2 | n.p. |

**Table 2.** Fold-change of amino acid content of sulfonylurea-treated tissues compared to control. Fold-change presented if significant (two-sample equal variance t-test for apple and flowering quince, paired t-test for banana; one-tailed test for valine, leucine and isoleucine; two-tailed test for all others; α = 0.05). Branched-chain amino acids highlighted in red. Proline was not determined in apple and ornamental quince samples. For quantified amino acid levels and p-values, see Supplemental Tables.

|  | apple peel |  |  | banana pulp | flowering quince peel |
| --- | --- | --- | --- | --- | --- |
| amino acid | 'Gala' | 'Empire' | 'Jonagold' | 'Valery' | 'Dr. Banks Pink' |
| alanine | ▲ 2.7 | — | — | ▲ 4.0 | — |
| arginine | — | — | — | — | — |
| asparagine | — | — | — | — | — |
| aspartate | ▲ 2.3 | — | — | ▲ 3.3 | — |
| cysteine | — | ▲ 2.6 | ▲ 4.1 | ▲ 2.5 | — |
| glutamate | ▲ 1.5 | — | — | ▲ 2.4 | — |
| glutamine | ▲ 2.5 | — | — | — | — |
| glycine | — | — | — | ▲ 1.6 | — |
| histidine | — | — | — | — | ▲ 1.7 |
| *isoleucine* | ▼ 6.1 | ▼ 30.2 | ▼ 24.1 | — | ▼ 1.9 |
| *leucine* | — | — | — | ▼ 1.5 | — |
| lysine | — | — | — | — | — |
| methionine | — | ▲ 2.1 | ▲ 2.2 | — | ▲ 2.5 |
| phenylalanine | — | — | — | — | — |
| proline | n.d. | n.d. | n.d. | ▲ 1.9 | n.d. |
| serine | ▲ 3.0 | — | ▲ 1.9 | ▲ 1.9 | ▲ 3.0 |
| threonine | ▲ 1.8 | — | — | — | — |
| tryptophan | — | — | — | — | — |
| tyrosine | — | — | — | ▲ 1.4 | — |
| *valine* | — | — | — | ▼ 7.0 | ▼ 3.6 |
| total | ▲ 1.9 | — | — | ▲ 1.5 | — |

While treatment of rimsulfuron had no discernable effect on butyl acetate, concentrations of pentyl acetate were more abundant in all three of the cultivars when treated (**Table 1; Supplemental Fig. S1; Supplemental Fig. S2; Supplemental Table S1**). Ethyl acetate was slightly lower in ‘Gala’ and ‘Empire’ fruits. Propyl acetate and hexyl acetate were marginally reduced in ‘Empire’ and ‘Jonagold’ headspaces, respectively.

#### Amino acids

Isoleucine was found to be significantly reduced in the peels of all three cultivars after rimsulfuron treatment. Interestingly, neither valine nor leucine, which also depend upon AHAS for synthesis, were found to be different in the treated tissues (**Table 2; Supplemental Table 2; Supplemental Table 3; Supplemental Table 4**).

Notably no other amino acid was reduced following treatment, however while several were found to be somewhat elevated, no patten of change was discerned across the cultivars. Only ‘Gala’ fruit had a significant increase of total free amino acid content after treatment.

#### α-Keto acid feedings

To discern if herbicide-treated fruit were still capable of aroma production, and to test the importance of substrate availability, branched-chain ⍺-ketoacids were fed, with and without inhibitor, to ‘Empire’ and ‘Jonagold’ peel tissues.

Exogenously fed ⍺-keto-β-methylvalerate, the ⍺-ketoacid of isoleucine, led to a partial rescue of all 2-methylbutyl and 2-methylbutanoate esters of the rimsulfuron-treated fruit, indicating ester synthesis capability was sustained in herbicide-treated fruit and that reduced volatile levels were a result of precursor scarcity, not herbicide toxicity (**Supplemental Fig. S1; Supplemental Fig. S2**).

Both cultivars were also capable of converting exogenous ⍺-ketoisovalerate and ⍺-ketoisocaproate, the ⍺-ketoacids of valine and leucine, respectively, into an abundance of their respective iso-branched-chain aldehydes, alcohols, alkyl ester elements (those derived from alcohols) and alkanoate ester elements (those derived from acyl-CoAs), none of which are normally produced in appreciable quantities by apple fruit, but are abundant in banana (**Fig. 3B; Supplemental Fig. S1; Supplemental Fig. S2; Supplemental Table S5**). Furthermore, activity of isopropylmalate synthase, the enzyme that facilitates extension of ⍺-ketoisovalerate to ⍺-ketoisocaproate, was evident by the observed emanation of 3-methylbutanal, 3-methylbutanol, and several 3-methylbutyl esters from fruit fed with ⍺-ketoisovalerate (**Fig. 2**; **Fig. 3B; Supplemental Table S5)**.

In general, the fruit preferentially converted the supplied ⍺-ketoacids into alkyl ester elements rather than alkanoate ones (**Supplemental Fig. S1; Supplemental Fig. S2**). Furthermore, while each of the exogenously supplied ⍺-ketoacids was converted into similar quantities of alkyl ester elements by each cultivar, ⍺-ketoisocaproate was metabolized into substantially less ethyl 3-methylbutanoate relative to the conversion of ⍺-keto-β-methylvalerate and ⍺-ketoisovalerate to ethyl 2-methylbutanoate and ethyl 2-methylpropanoate, respectively, suggesting nuance within the intermediary enzymatic steps between ⍺-ketoacid and volatile (**Fig. 3B; Supplemental Fig. S1; Supplemental Fig. S2**).

#### Acetate feeding

Our ability to inhibit AHAS activity provided a valuable opportunity to investigate the interplay of the citramalate synthase pathway and branched-chain amino acid metabolism in apple fruit (**Fig. 2**). Citramalate synthase facilitates the synthesis of ⍺-ketobutyrate, one of the substrates of AHAS for the eventual production of isoleucine (Sugimoto, et al., 2021). While threonine deaminase can also produce ⍺-ketobutyrate, the citramalate synthase pathway is believed to be the principal source in ripening apple peel as citramalate synthase, unlike threonine deaminase, is not allosterically feedback inhibited by isoleucine. Furthermore, citramalate synthase, whose primary substrates are pyruvate and acetyl-CoA, has been observed to be capable of also facilitating the extension of ⍺-ketobutyrate to ⍺-ketovalerate, as well as ⍺-ketovalerate to ⍺-ketocaproate, via further incorporation of acetyl-CoAs. These straight-chain ⍺-ketoacids are hypothesized to be potential precursors to straight-chain esters via analogous means as branched-chain ⍺-ketoacids and branched-chain esters. Thus, the citramalate synthase pathway of apple fruit not only contributes to branched-chain ester biosynthesis, but to that of straight-chain esters as well. However other means of straight-chain ester biosynthesis, such as those facilitated by the lipoxygenase and β-oxidation pathways, are thought to supply the principal proportion of straight-chain ester precursors. Nonetheless, by inhibiting AHAS, ⍺-ketobutyrate cannot be metabolized to isoleucine, effectively shunting flux of the pathway towards further straight-chain elongation. To explore the potential shifts of metabolism, we fed 1,2-^13^C_2_ acetate, which is readily metabolized to acetyl-CoA, with and without inhibitor, to track possible shifts of carbon flux in the fruit.

Attempts were made to feed the labeled acetate to the three cultivars, however incorporation by ‘Gala’ fruit was very poor as compared to ‘Empire’ and ‘Jonagold’ fruits, thus only the results of these latter cultivars were considered (**Supplemental Fig. S3; Supplemental Fig. S4; Supplemental Fig. S5**). Overall, the patterns of enrichment of control ‘Jonagold’ fruit were similar to that previously observed, however ‘Empire’ fruit did detract slightly from ‘Jonagold’ patterning (Sugimoto, et al., 2021).

Because of the herbicide inhibition of AHAS, rimsulfuron-treated ‘Empire’ produced very low amounts of methyl 2-methylbutanoate. No isotopolog enrichment patterning was observed. The production of methyl 2-methylbutanoate by rimsulfuron-treated ‘Jonagold’ was below the limits of detection, so enrichment could not be assessed.

No significant differences of ^13^C incorporation were found in rimsulfuron-treated fruits of either cultivar for methyl acetate, methyl propanoate or methyl butanoate. However, rimsulfuron treatment resulted in an enrichment of the M+1 and M+2 methyl pentanoate isotopologs in both cultivars. These enrichments were complemented with a corresponding significant decrease of the M isotopologs. Lastly, the M+1 isotopolog fraction of methyl hexanoate was enriched in rimsulfuron-treated ‘Jonagold’ fruit.

#### Sensory discrimination and population genetics

Our ability to inhibit 2-methylbutyl and 2-methylbutanoate ester biosynthesis, and the inodorous, non-volatile nature of the herbicide provided an interesting opportunity to investigate the sensory impact of these compounds. To determine if humans can detect the absence of branched-chain volatiles in apple fruits, we presented panelists with slices of rimsulfuron-treated or untreated ‘Jonagold’ peel in a duo-trio discrimination test. In a duo-trio test, panelists are instructed to select which one of two blinded samples (untreated, treated) matches a reference sample (untreated). Panelists were able to discriminate between samples that included or lacked these compounds, correctly matching the untreated fruit to the reference significantly more often than by chance (**Fig. 4A; Supplemental Table S6**; n = 240; 64% correct; p = 6.7*×*10^-6^). These results inspired us to further investigate the implications this may have had on apple breeding and propagation.

**Figure 4.**
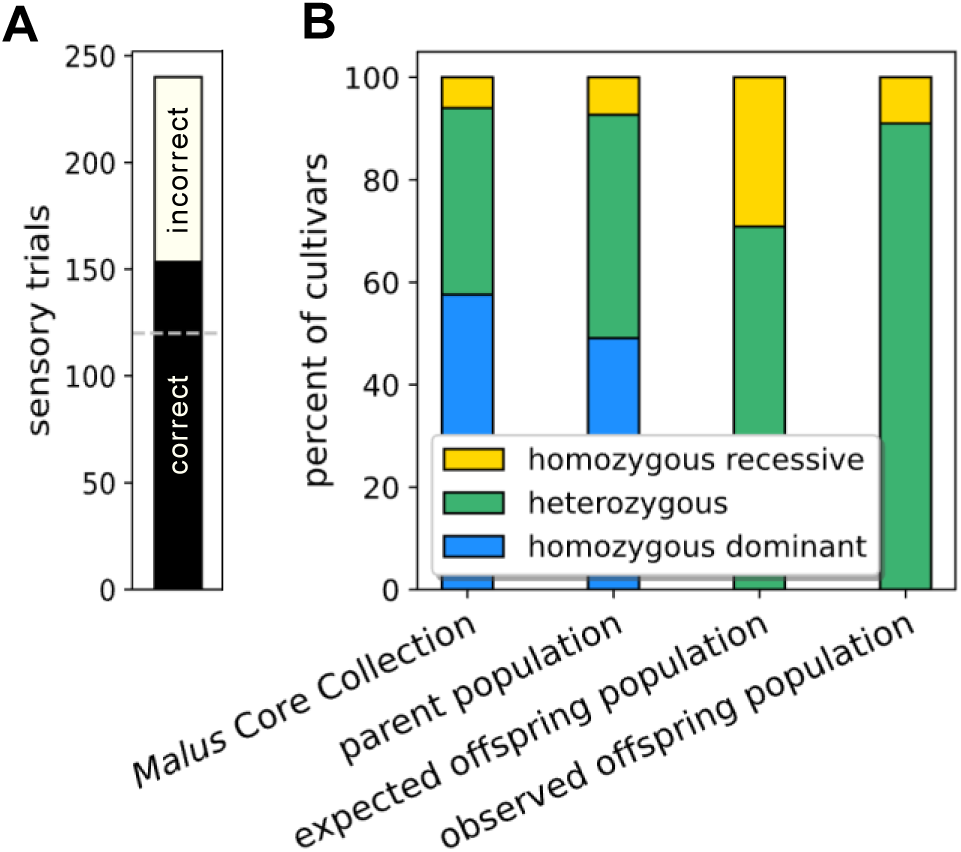
Results of sensory trials and population genetics analyses. A) Compiled results from duo-trio discrimination tests between rimsulfuron and water treated ‘Jonagold’ fruit. Dashed line shows expected number of correct trials if determined by chance. (n = 240, 64% correct, *p* = 6.7×10^-6^). B) Distributions of citramalate synthase genotypes. *Malus* Core Collection is distribution of 99 cultivars as assessed by Sugimoto et al., 2021. Parent population (n = 55) is derived from USDA Geneva *Malus* Core Collection to mimic population definitions of Migicovsky et al., 2022 through the recusal of recent breeding line hybrids and represents the population genetics of potential mates with ‘Cox’s Orange Pippin’. Offspring populations (n = 32) are derived from ‘Cox’s Orange Pippin’ (homozygous recessive) × parent population as identified by Migicovsky et al., 2022. Expected offspring population vs observed offspring population (χ^2^, *p* = 0.0141). See Supplemental Text for further detail.

When exogenous herbicide application is not considered, the ability of an apple cultivar to produce 2-methylbutyl and 2-methylbutanoate esters is determined by a dominant-recessive trait where synthesis is dependent upon the presence of at least one active allele of citramalate synthase (Sugimoto, et al., 2021). Among cultivars of the USDA Geneva *Malus* 4-Copy Core Collection previously analyzed, only 6.1% were homozygous recessive, despite 42.4% of cultivars carrying the recessive allele (Sugimoto, et al., 2021). Homozygous recessive cultivars likewise had significantly lower ratios of 2-methylbutyl and 2-methylbutanoate esters to straight-chain esters as compared to cultivars with at least one copy of the active allele (Sugimoto, et al., 2015). While this data demonstrates that cultivars lacking 2-methylbutyl and 2-methylbutanoate esters are uncommon, it cannot be determined if such a distribution is a result of natural or artificial selection.

However, among the cultivars screened, ‘Cox’s Orange Pippin’, which has been identified through pedigree and sequencing-based analyses as a common breeding parent to many cultivars (Muranty, et al., 2020) (Noiton & Alspach, 1996), was found to be homozygous recessive for citramalate synthase (Sugimoto, et al., 2021). If there were no selection pressure upon citramalate synthase activity, then one would expect a relatively large proportion of the offspring of ‘Cox’s Orange Pippin’ to likewise be homozygous recessive and thus have much reduced production of 2-methylbutyl and 2-methylbutanoate esters.

To investigate the population genetics of citramalate synthase, and thus branched-chain esters in apples, we sequenced the consequential single nucleotide polymorphism of citramalate synthase from thirty-two cultivars within the USDA Geneva *Malus* Core Collection that have been previously identified as being derived from ‘Cox’s Orange Pippin’ (Migicovsky, et al., 2021; Muranty, et al., 2020; Howard, et al., 2023). Given our population definitions (see Supplemental Text), we would expect 29.1% of ‘Cox’s Orange Pippin’s offspring to be homozygous recessive if no selection is occurring, however, only three cultivars (9.4%) were homozygous recessive (**Fig. 4B**; **Supplemental Table S7** χ^2^; p = 0.0141; see Supplemental Text for further details).

### Banana

The application of halosulfuron, another sulfonylurea AHAS inhibitor, led to a reduction of the headspace concentrations of every iso-branched-chain ester and alcohol analyzed in ‘Valery’ banana fruits (**Fig. 3C**; **Table 1; Supplemental Table S8**). Surprisingly, ethyl acetate, butyl acetate, and butyl butanoate were also less in treated tissues. None of the sec-branched esters or their related volatiles (2-pentanone, 2-pentanol, 1-methylbutyl acetate, and 1-methylbutyl butanoate) were affected by halosulfuron treatment.

Analogous to our results in apple fruit, application of exogenous branched-chain ⍺-ketoacids to tissues treated with inhibitor led to a recovery of the iso-branched-chain esters (**Supplemental Fig. S6**). Furthermore, banana fruit were capable of metabolizing the ⍺-ketoacid of isoleucine, ⍺-keto-β-methylvalerate, into a variety of anteiso-branched-chain volatiles, as has been previously observed (**Supplemental Fig. S6; Supplemental Table S9**) (Wyllie & Fellman, 2000). While none of the applied ⍺-ketoacids led to a recovery of ethyl acetate or butyl acetate in this reported trial, we would be remiss to fail to mention that during method testing we did occasionally observe a marked recovery of ethyl acetate and butyl acetate after ⍺-ketoisocaproate application, however further investigation is needed to better understand these observations.

Valine and leucine were significantly less abundant in halosulfuron-treated fruit, whereas isoleucine levels were unchanged (**Table 2; Supplemental Table S10**). No other amino acids were reduced in halosulfuron-treated fruits. Several amino acids were found to be higher after herbicide treatment with the total content of free amino acids having increased.

The observed independence of sec-branched-chain volatile biosynthesis to AHAS activity, as well as the unknown nature of their biochemical origins, inspired us to further explore this understudied component of banana fruit aroma. To investigate 2-pentanone and 2-pentanol and their possible role as precursors to 1-methylbutyl esters, as well as the importance of substrate availability to aroma biosynthesis in bananas, sections of ‘Horn Plantain’ pulp were fed 2-pentanone, 2-pentanol, or butanol to assess this cultivar’s ability to metabolize these compounds into esters.

‘Horn Plantain’ fruit have, compared to ‘Valery’, a very simple aroma profile that is restricted to ethyl acetate, 2-methylpropyl acetate, and 3-methylbutyl acetate. Notably, the cultivar produces very little butyl acetate or 1-methylbutyl acetate and has very limited production of butanoate esters.

The feeding of 2-pentanol, but not 2-pentanone, resulted in the accumulation of 1-methylbutyl acetate in the headspace (**Fig 3D; Supplemental Fig. S7**). Furthermore, 2-pentanol feeding did not lead to an increase of 2-pentanone or vice versa. Butyl acetate and 2-methylpropyl butanoate production was increased when the tissues were fed butanol, however the resulting level of 2-methylpropyl butanoate (∼0.6 nmol ᐧ L^-1^) was much lower than that of butyl acetate (168.0 nmol ᐧ L^-1^). No other esters were determined to be affected by the feedings.

### Flowering quince

To demonstrate the reach of our hypothesis that *de novo* synthesis is a major contributor to branched-chain ester biosynthesis in ripening fruits, we applied rimsulfuron to the highly aromatic fruit of an ornamental quince hybrid (*Chaenomeles* ×*superba*, cv. Dr. Banks Pink; **Supplemental Fig. S8**); a fruit, when compared to apples, whose means of aroma biochemistry are essentially uninvestigated.

The aroma profile of the small, dense fruit was found to be dominated by the terpene linalool and the phenylpropene estragole, but low levels of the straight-chain esters propyl acetate, butyl acetate and ethyl butanoate, as well as the branched-chain esters 2-methylpropyl acetate and 2-methylbutyl acetate are present (**Table 1; Supplemental Table S11**). As this species is a member of Maleae (the tribe of Rosaceae that includes apples and pears), it was assumed that the peel is the site of aroma biogenesis (Guadagni, et al., 1971). Thus, rimsulfuron was applied to the peels of aroma-active fruits and the effect upon aroma production and amino acid content was analyzed.

Rimsulfuron-treated fruits had less 2-methylpropyl acetate and 2-methylbutyl acetate, but more propyl acetate than untreated fruits. No difference was observed for butyl acetate, ethyl butanoate, estragole, or linalool (**Table 1; Supplemental Table S11**).

Valine and isoleucine were reduced in rimsulfuron-treated peel tissue (**Table 2; Supplemental Table S12**), but no difference was seen for leucine, nor was the concentration of any other amino acid diminished by herbicide treatment. While histidine, methionine, and serine underwent modest increases, the total free amino acid pool level was not significantly altered by treatment.

## Discussion

The categorical suppression of every isoleucine-, valine-, and leucine-related volatile analyzed in the fruit tissues treated with an AHAS inhibitor emphatically demonstrates that these fruit tissues rely heavily, or perhaps solely, upon newly synthesized precursors for the production of these important sensory compounds. If the substrates for branched-chain volatile biosynthesis are procured through an alternative pathway, such as protein degradation, then application of the inhibitors should have had no effect upon volatile production based on our current understanding of their mode of action (Shaner & Reider, 1986). However the volatiles in question, as well as their related amino acids, were in fact reduced by the inhibitors, thus demonstrating that past observations of the marked increases of only branched-chain amino acids that have related volatiles being produced, i.e. isoleucine in apple, or valine and leucine in banana, are not a product of some coincidental catabolic phenomenon, but are due to deliberate enhancements of specific portions of these branched-chain pathways (Sugimoto, et al., 2011) (Sugimoto, et al., 2015) (Alsmairat, et al., 2018).

Theoretically, if branched-chain esters are directly derived from branched-chain ⍺-ketoacids, as is widely speculated, then interconversion of the amino acids and ⍺-ketoacids may be unnecessary. Thus, the accumulated branched-chain amino acids in ripening fruits are a byproduct of, *and not precursors nor intermediates to,* branched-chain volatile biosynthesis. It would thus be much more accurate to refer to branched-chain volatiles as being ‘related to’ branched-chain amino acids, as opposed to being ‘derived from’ them.

The use of sulfonylureas as a biochemical tool also allowed for extensive experimentation of the nuances of the various biochemical pathways interconnected with AHAS. The subtle preference of apple fruits to convert branched-chain ⍺-ketoacids into alkyl ester elements, as well as the greater ability for ⍺-keto-β-methylvalerate and ⍺-ketoisovalerate to be converted into alkanoate ester elements relative to ⍺-ketoisocaproate can be more easily observed when *de novo* influx of these substrates is inhibited, giving greater control over experimental conditions. Additionally, as the herbicide application led to no apparent changes of ripening behavior, application likewise allowed for the comparison of fruit with select modification of the aroma profiles, as was utilized for our sensory analysis.

Our results likewise demonstrate that apple and banana fruit, despite not normally producing appreciable amounts of all three classes of anteiso- and iso-branched-chain esters, have no general hindrance to synthesizing said esters when a supply of the corresponding ⍺-ketoacids is present. Together these experiments, in conjugation with the ability of ‘Horn Plantain’ fruit to incorporate 2-pentanol and butanol into their respective esters, highlight the importance of substrate availability to ester biosynthesis. This is in opposition to the notion that the substrate preference of alcohol acyl transferase, the enzyme that reacts alcohols and acyl-CoAs to form esters, is paramount (Beekwilder, et al., 2004) (Aharoni, et al., 2000).

Sulfonylurea application also allowed for further investigation of the citramalate synthase pathway in apple. The unanimous increase of pentyl acetate content as well as methyl pentanoate ^13^C enrichment in treated apple fruits suggests that a substantial proportion of pentyl and pentanoate esters are dependent upon the citramalate synthase pathway. It also suggests that the activity of AHAS draws flux away from the straight-chain elongation portion of the citramalate synthase pathway. Our inability to detect consistent differences of the other straight-chain esters (e.g., propyl, propanoate, butyl, and butanoate) may have been due to dilution of the substrate pool due to influx from other sources, such as the lipoxygenase and β-oxidation pathways.

Furthermore, by inhibiting synthesis of 2-methylbutyl and 2-methylbutanoate esters we gained the opportunity to investigate not only the biosynthesis of these compounds, but also their sensory impact. The ability of humans to perceive the absence of these compounds, despite the relative complexity of an apple fruit’s aroma profile, is impressive when one considers that differing aroma profiles are increasingly difficult to distinguish as they become more similar (Bushdid, et al., 2014). It thus seems feasible that, whether consciously or not, humans have been selecting for apple cultivars that produce these branched-chain compounds, highlighting their importance to humankind.

In banana fruit, the use of these inhibitors has allowed for the suggestion of a previously unknown means of butyl and butanoate ester biosynthesis that appears to be unique to banana fruit since similar results were not observed in apple or flowering quince. Apart from some specialized instances of straight-chain 1-C elongation from α-ketobutyrate that have been documented in apple [via citramalate synthase (Sugimoto, et al., 2021)] and *Solanaceae* (Kroumova & Wagner, 2003), it has largely been assumed that butyl and butanoate esters are derived from β-oxidation of fatty acids; a catabolic process. As treatment of tissues with AHAS inhibitors does not lead to inhibition of fatty acid synthesis (Shaner & Singh, 1997), our results suggest that the source of butyl and butanoate esters in banana fruit is originally from branched-chain amino acid metabolism.

The synthetic mechanism for butyl and butanoate ester formation in banana is unclear. However, some of the first scientific work that attempted to establish a relationship between leucine and 3-methylbutyl esters also found that bananas fed U-^14^C-leucine produced a significantly enriched volatile fraction containing butyl butanoate and 1-methylbutyl butanoate (Tressl & Drawert, 1973). In light of these results, we propose that ⍺-ketoisocaproate is the metabolite that banana fruit use to bridge branched-chain amino acid and butanoate metabolisms, however future work elucidating how this is facilitated is needed to confirm this notion. ‘Horn Plantain’ fruit, which produce copious amounts of iso-branched-chain esters, and yet showed no barrier to incorporation of butanol into butyl esters, may be an excellent tool to help discern this pathway.

The sulfonylurea-induced inhibition of ethyl acetate synthesis in banana fruit is surprising given there is no described link between the paths of branched-chain α-ketoacid formation and the synthesis of either ethanol or acetate. However another past study, likewise feeding U-^14^C-leucine to ripening banana fruit, found enrichment of both the alkyl and alkanoate portions of 3-methylbutyl acetate (Myers, et al., 1970), leading to the suggestion that even the precursors of ethyl acetate may be potentially derived from branched-chain amino acid metabolism in banana fruit.

The lack of suppression of 2-pentanol, 2-pentanone, and 1-methylbutyl esters by the sulfonylurea herbicide strongly suggests that these sec-branched-chain compounds of banana fruit are derived from a source that is not within the sphere of AHAS’s influence. Furthermore, the ability of ‘Horn Plantain’ fruit to incorporate 2-pentanol, but not 2-pentanone, into 1-methylbutyl esters hints that the more stable ketone, detected in natural emanations from ‘Valery’, may be a dead-end byproduct of 2-pentanol oxidation. While we did not observe such an interconversion, it is possible that the short incubation period provided (1-2 hours) was not sufficient. It is also striking that banana fruit should produce multiple forms of branched-chain esters (iso- and sec-branched), that are of seemingly wholly independent origins.

Lastly, our results with flowering quince demonstrate the meaningful strides in dissecting the paths of branched-chain ester formation made possible when utilizing the targeted and well-described effectiveness of herbicides. Principally, our results for flowering quince indicate that these fruit, like apple and banana fruit, rely upon *de novo* synthesis to produce branched-chain volatiles. However, while our results cannot indicate whether the citramalate synthase pathway is present in flowering quince, the sole increase of propyl acetate after sulfonylurea application is, perhaps, telling. Propyl acetate is one of the ester products of ⍺-ketobutyrate following its decarboxylation. However, given the lack of enhancement of longer straight-chain products, as observed in apple fruit, suggests 1-C elongation of α-ketoacids is likely not occurring in this fruit.

Collectively, our results, which rely upon findings using fruit as diverse as being from a perennial deciduous dicot tree and an herbaceous tropical monocot, reject the hypothesis that branched-chain esters are derived from preexisting branched-chain amino acids and instead support the hypothesis that these esters are the product of *de novo* precursor biosynthesis.

## Conclusion

In this work, a powerful tool for the study of volatile biochemistry has been described. Interestingly, we could find no similar published work utilizing herbicides to explore pathways for ester biosynthesis. Through the application of sulfonylurea herbicides, we have been able to demonstrate, among other things, the importance of *de novo* synthesis of branched-chain ⍺-ketoacids in the production of aroma compounds, highlight the importance of branched-chain compounds to the complex aroma profiles of apple fruits, gain insight into the nuances of ⍺-ketoacid metabolism for ester biosynthesis, and potentially uncover a previously unknown biochemical pathway of aroma biosynthesis. No doubt the future use of herbicides on other fruits, whether those used herein or others that target different metabolic networks, will continue to shed light on the importance of substrate availability and aroma biochemistry in general.

Lastly, it is worthwhile to pursue the logical extension to our results. As we have demonstrated, apple and banana fruits produce anteiso- and iso-branched-chain esters from newly synthesized precursors. However, the pathways that produce said precursors are normally regulated by strict allosteric feedback mechanisms. The ability of these fruits to increase flux through these pathways during ripening seems paradoxical. While citramalate synthase explains how apple fruit circumvent such regulation, it is still unknown what regulatory changes occur in ripening banana fruits.

## Materials and methods

### Plant material

‘Gala’, ‘Empire’, and ‘Jonagold’ apple (*Malus* ×*domestica* Borkh.) fruit were harvested from local orchards at commercial maturity and transported to the laboratory during the 2022 season. Developmentally, the fruit were harvested at the onset of ripening, but no aroma could be discerned subjectively. ‘Gala’ and ‘Empire’ fruits began treatment (see below) immediately after arrival. ‘Jonagold’ fruit were held in air at 0 ℃ for two days before transfer to 1.5 O_2_, 3% CO_2_, 0 ℃. ‘Jonagold’ fruit were held in these conditions for twelve days before the initiation of treatment.

Mature, green banana (*Musa* spp. AAA group, Cavendish subgroup, cv. Valery; *Musa* spp. AAB group, Plantain subgroup, cv. Horn Plantain) fruit that had not been treated with ethylene were obtained from a local supermarket produce distribution and ripening center (Meijer/Chiquita, Lansing, MI). ‘Horn Plantain’ fruit were held at room temperature (22 ℃) for 2-3 days before treatment. ‘Valery’ fruit were held at 13.5 ℃ and 95% RH until treatment, which typically took place in the first two weeks of holding.

‘Dr. Banks Pink’ flowering quince (*Chaenomeles* ×*superba*) fruit were collected from accession CC7985*05 on the campus grounds of Michigan State University. Fruits were actively producing aroma at the time of harvest.

### Treatment

Whole apple and quince fruit were treated daily with freshly made herbicide or water solution (1 mM rimsulfuron, made from DuPont™ Matrix® SG (25% w/w active ingredient), 0.1% Tween 20) before preparation with further treatments. Application was by placing 3 mL of the herbicide solution on the fruit and plastic gloves and rubbing the surface of the fruit until the entire surface was wetted. Whole fruit routinely had their headspace volatiles sampled for aroma production and treatment efficacy to determine appropriate times for further treatments.

Acetate and ⍺-ketoacid feedings were performed on excised peel discs of apple peel as previously described (Sugimoto, et al., 2021). See Supplemental **Methods** for further detail.

Banana fruit were prepared while mature and green as either ∼1 cm^2^ square sections of pulp or 5 mm transverse cross-sections of whole fruit as inspired by (Palmer & McGlasson, 1969). Samples were treated with 0.1% Tween 20 with or without herbicide (0.5 mM halosulfuron, made from Sandea® (75% w/w active ingredient)), incubated with ethylene to induce ripening, and then analyzed several days later when ripe. Temperature was 22 °C throughout the handling process. See **Supplemental Methods** for further detail.

### Volatile analysis

The volatiles produced from the samples were analyzed using a solid-phase micro extraction (SPME) fiber (65 μm PDMS-DVB; Supelco Analytical, Bellefonte, PA) and a gas chromatograph (GC; HP-6890, Hewlett-Packard, Wilmington, DE) coupled to a time of flight mass spectrometer (MS; Pegasus III, LECO, St. Joseph, MI). Treated excised apple peel discs and banana pulp sections were incubated in 22-mL glass vials overnight prior to sampling. Whole quince fruit and banana slices were incubated for 20 min in 1-L sealed Teflon jars before sampling. Compounds were identified by comparison with the retention time and mass spectrum of authenticated reference standards and spectra (National Institute of Standards and Technology Mass Spectral Search Program Version 2.0, 2001). Volatiles were quantified by calibration with a standard of 59 authenticated compounds (Sigma-Aldrich Co., St. Louis, MO and Fluka Chemika, Seelza, Germany). After volatile analysis, apple peel disks, collected peel of quince, and ‘Valery’ pulp section samples were held at -80 ℃ for amino acid analysis. See Supplemental Methods for further detail.

### Sensory evaluation

Human subjects (n = 20) with a normal sense of smell were recruited from the greater Lansing, MI area to participate in a sensory study on fruit aroma. Participants came into the Sensory Lab for a single session during which they performed twelve duo-trio discrimination trials (n = 240). In each trial, subjects were presented with 3 sample vials: an untreated sample labeled “Reference”, a blinded untreated sample, and a blinded treated sample. Subjects were instructed to smell the reference and the two blinded vials and to select which vial matched the reference. ‘Jonagold’ fruit, previously held in controlled atmosphere storage as described above, were treated with herbicide or water solution (1 mM rimsulfuron, made from Matrix®, 0.1% Tween 20) as described above. Fruits were screened via GCMS, described above, to ensure successful treatment and suppression of 2-methylbutyl and 2-methylbutanoate esters. The day before the sensory study, 1 x 5 cm segments weighing 2.5 – 3 g, were prepared from the apples. Cortex tissue was trimmed to maximize the proportion of peel tissue. Fruit segments were then placed in 40-mL amber vials and sealed with PTFE-lined caps. Samples were allowed to incubate at least 2 hours before the first sensory evaluation session. This protocol was reviewed and determined to qualify for exempt status by the Michigan State University Institutional Review Board (Study 00008470). See Supplemental Methods for further detail.

### Citramalate synthase sequencing

The nonsynonymous SNP of the 387^th^ codon of citramalate synthase that has been found to be responsible for activity/inactivity of the enzyme (Sugimoto, et al., 2021) was Sanger sequenced at the Research Technology Support Facility Genomics Core at Michigan State University of cultivars previously identified as being first-degree relatives of ‘Cox’s Orange Pippin’ (Migicovsky, et al., 2021). See Supplemental Methods and Text for further detail.

### Amino acid analysis

Frozen samples were ground to a powder in liquid nitrogen-chilled mortar and pestles. Tissues were extracted with 1:1:1 water:acetonitrile:ethanol spiked with U-^13^C,^15^N labeled amino acids (MilliporeSigma) before filtering with 0.2 μm nylon centrifugal filter units. Filtrate was diluted with perfluorohexanoic acid spiked with internal standard prior to analysis with a Xevo TQ-S Micro UPLC (H-Class)-MS/MS (Waters, Milford, MA) at the Michigan State University Mass Spectrometry and Metabolomics Core. Samples were quantified against a likewise prepared serial dilution of amino acid stocks. See Supplemental Methods for further detail.

## Acknowledgments and Funding

P.E. greatly appreciates the kind contributions of banana fruit from Dan Gregory and of herbicide materials from Dr. Debalina Saha. R.B. acknowledges support from Michigan AgBioResearch and the U.S. Department of Agriculture National Institute of Food and Agriculture, Hatch project MICL002688. S.K.W. wishes to acknowledge Ryukoku University, Japan, for sabbatical leave support providing an Academic-Research Scholarship.

## Author Contributions

- Designed research: P.E., S.K.W., A.D., E.J.M., R.B.
- Performed research: P.E., S.K.W., A.D., E.J.M., R.B.
- Contributed new analytic/computation/etc. tools: P.E., S.K.W., R.B. (developed herbicide feeding techniques)
- Analyzed data: P.E., E.J.M., R.B.
- Wrote paper: P.E., R.B.

## Supplemental Figures

**Supplemental Figure S1.**
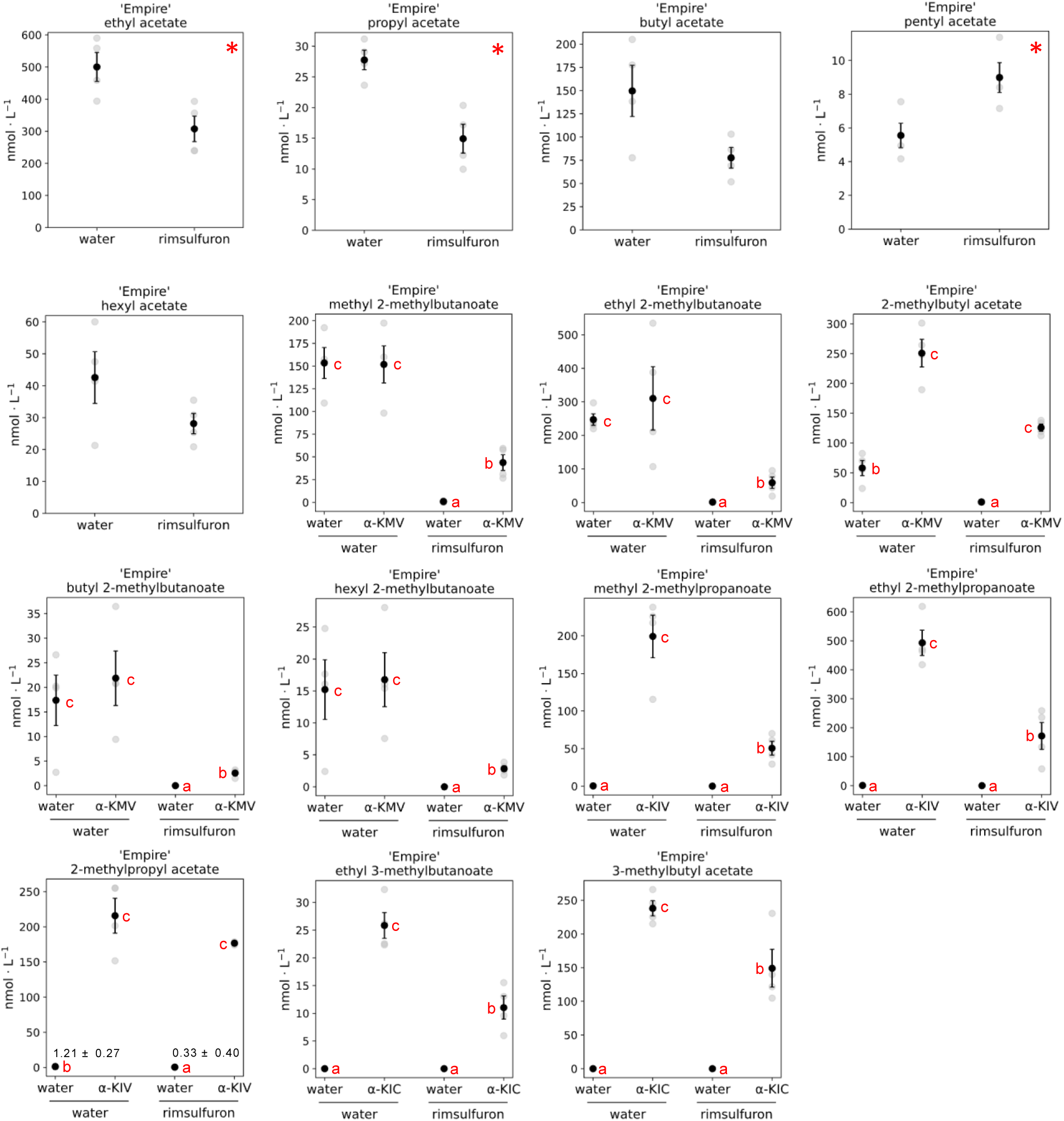
Volatile headspace concentrations of ‘Empire’ apple fruit peels treated with water or rimsulfuron and fed branched-chain ⍺-ketoacids. Presented as means ± ½ sᴅ of four biological reps. ⍺-KMV = ⍺-keto-β-methylvalerate; ⍺-KIV = ⍺-ketoisovalerate; ⍺-KIC = ⍺-ketoisocaproate. Significantly different straight-chain ester concentrations are denoted by * (two-tailed two-sample equal variance t-test, ⍺=0.05). Significantly different branched-chain ester concentrations are denoted by different letters adjacent to means (data transformed for statistical analysis via log(x+1) due to unequal variance of ⍺-ketoacid fed samples; Tukey’s test, ⍺=0.05). The concentrations of 2-methylpropyl acetate in tissues not fed with ⍺-KIV are shown in figure.

**Supplemental Figure S2.**
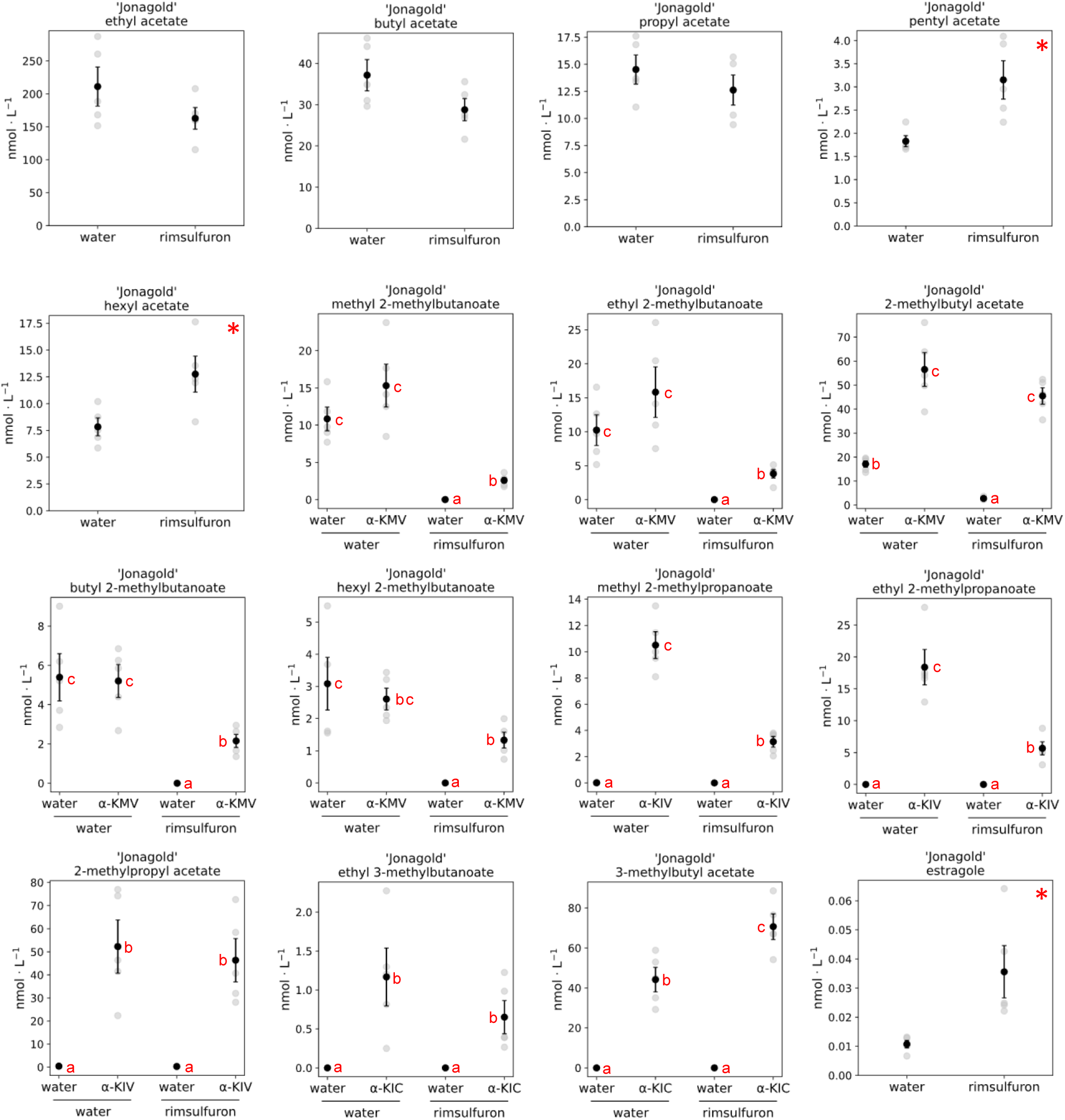
Volatile headspace concentrations of ‘Jonagold’ apple fruit peels treated with water or rimsulfuron and fed branched-chain ⍺-ketoacids. Presented as means ± ½ sᴅ of four biological reps. ⍺-KMV = ⍺-keto-β-methylvalerate; ⍺-KIV = ⍺-ketoisovalerate; ⍺-KIC = ⍺-ketoisocaproate. Significantly different straight-chain ester concentrations are denoted by * (two-tailed two-sample equal variance t-test, ⍺=0.05). Significantly different branched-chain ester concentrations are denoted by different letters adjacent to means (data transformed for statistical analysis via log(x+1) due to unequal variance of ⍺-ketoacid fed samples; Tukey’s test, ⍺=0.05).

**Supplemental Figure S3.**
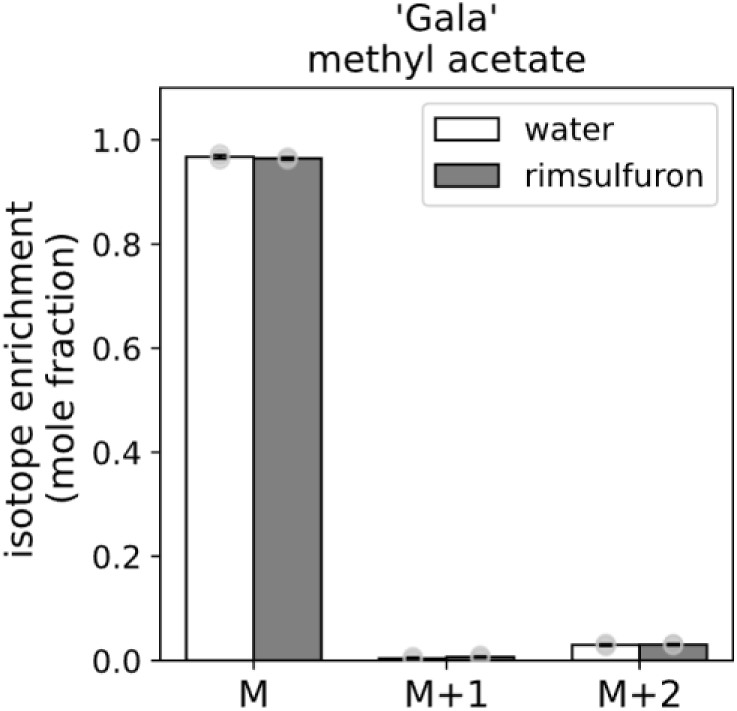
Mass isotopolog distribution of methyl acetate from ‘Gala’ apple fruit peels treated with water or rimsulfuron and fed 1,2- C_2_ acetate and methanol. Presented as means ± ½ sᴅ of two biological reps. Significantly different distributions are denoted by * (two-tailed two-sample equal variance t-test, ⍺=0.05).

**Supplemental Figure S4.**
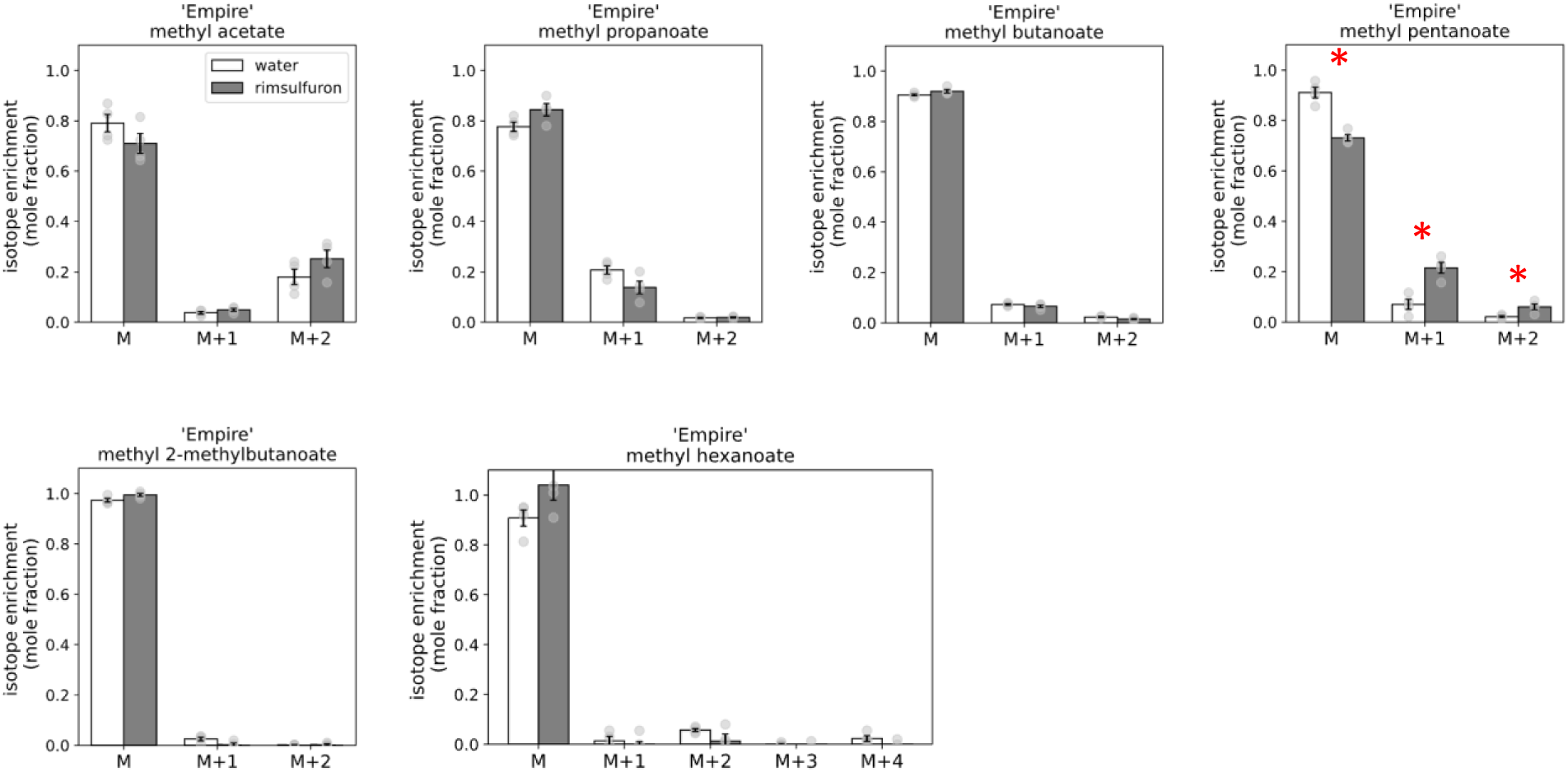
Mass isotopolog distribution of methyl esters from ‘Empire’ apple fruit peels treated with water or rimsulfuron and fed 1,2- C_2_ acetate and methanol. Presented as means ± ½ sᴅ of four biological reps. Significantly different distributions are denoted by * (two-tailed two-sample equal variance t-test, ⍺=0.05).

**Supplemental Figure S5.**
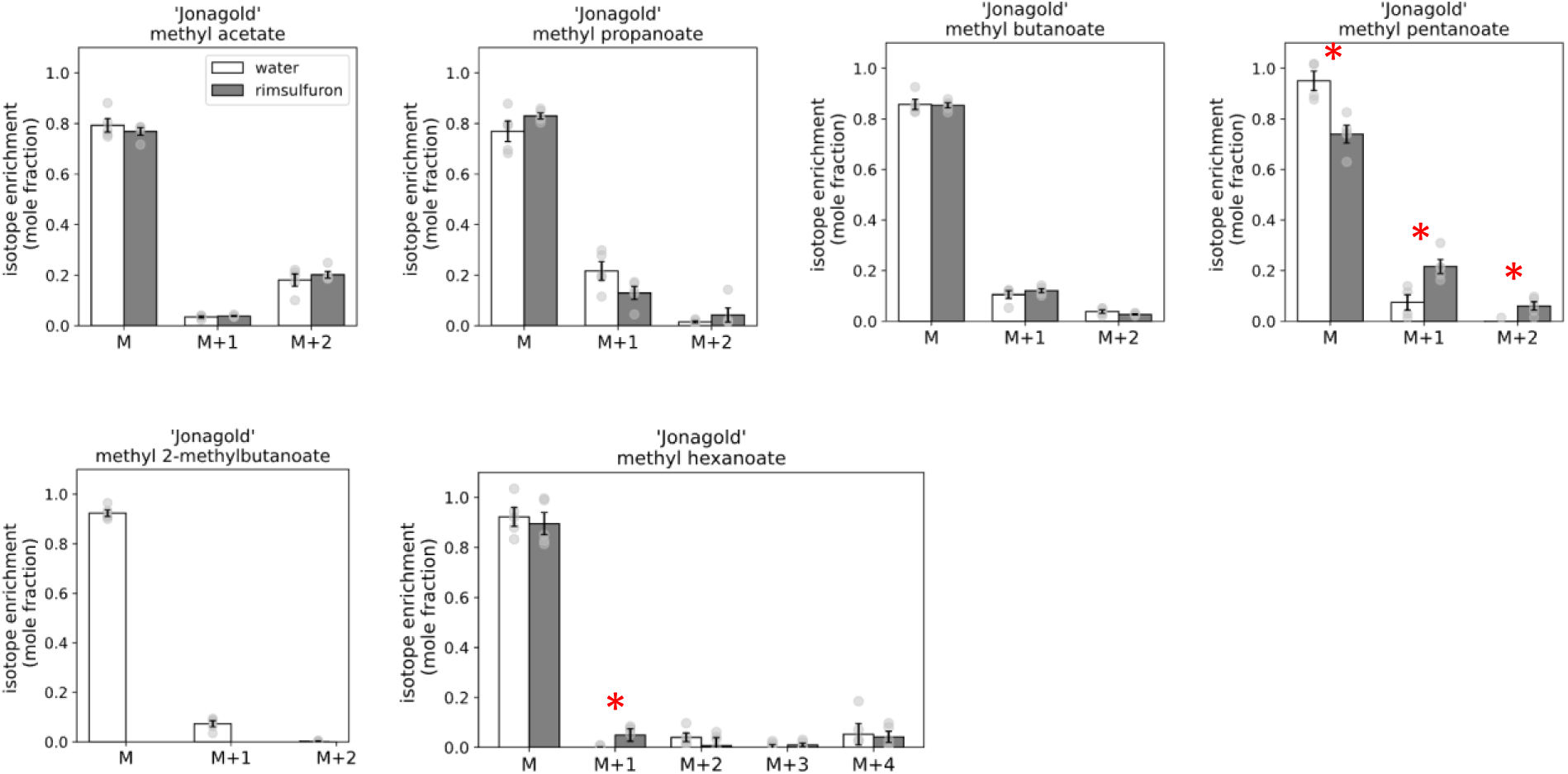
Mass isotopolog distribution of methyl esters from ‘Jonagold’ apple fruit peels treated with water or rimsulfuron and fed 1,2- C_2_ acetate and methanol. Presented as means ± ½ sᴅ of five biological reps. Significantly different distributions are denoted by * (two-tailed two-sample equal variance t-test, ⍺=0.05).

**Supplemental Figure S6.**
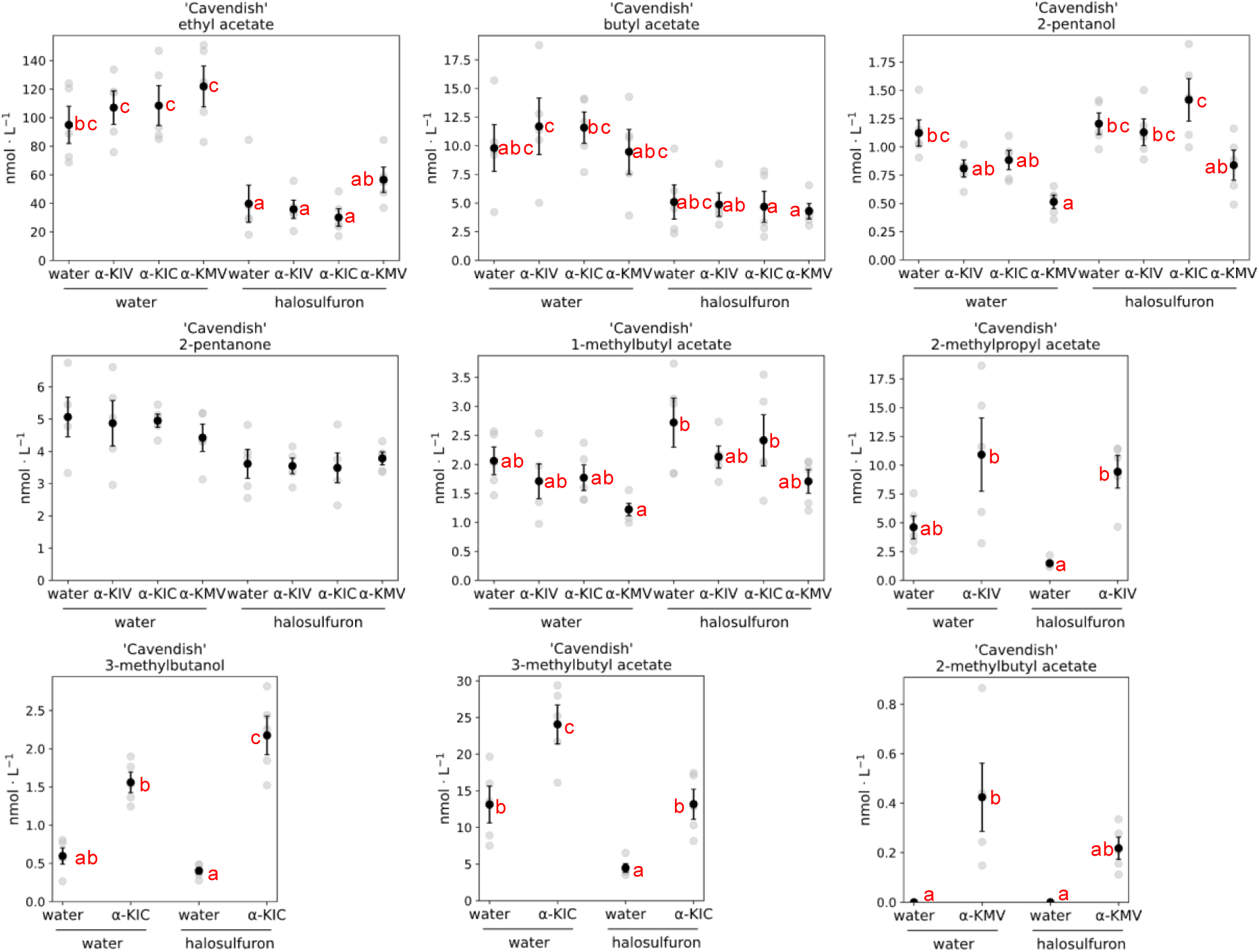
Volatile headspace concentrations of ‘Valery’ banana pulp sections treated with water or halosulfuron and fed branched-chain ⍺-ketoacids. Presented as means ± ½ sᴅ of five biological reps. ⍺-KMV = ⍺-keto-β-methylvalerate; ⍺-KIV = ⍺-ketoisovalerate; ⍺-KIC = ⍺-ketoisocaproate. Significantly different volatile are denoted by different letters adjacent to means (Tukey’s test, ⍺=0.05).

**Supplemental Figure S7.**
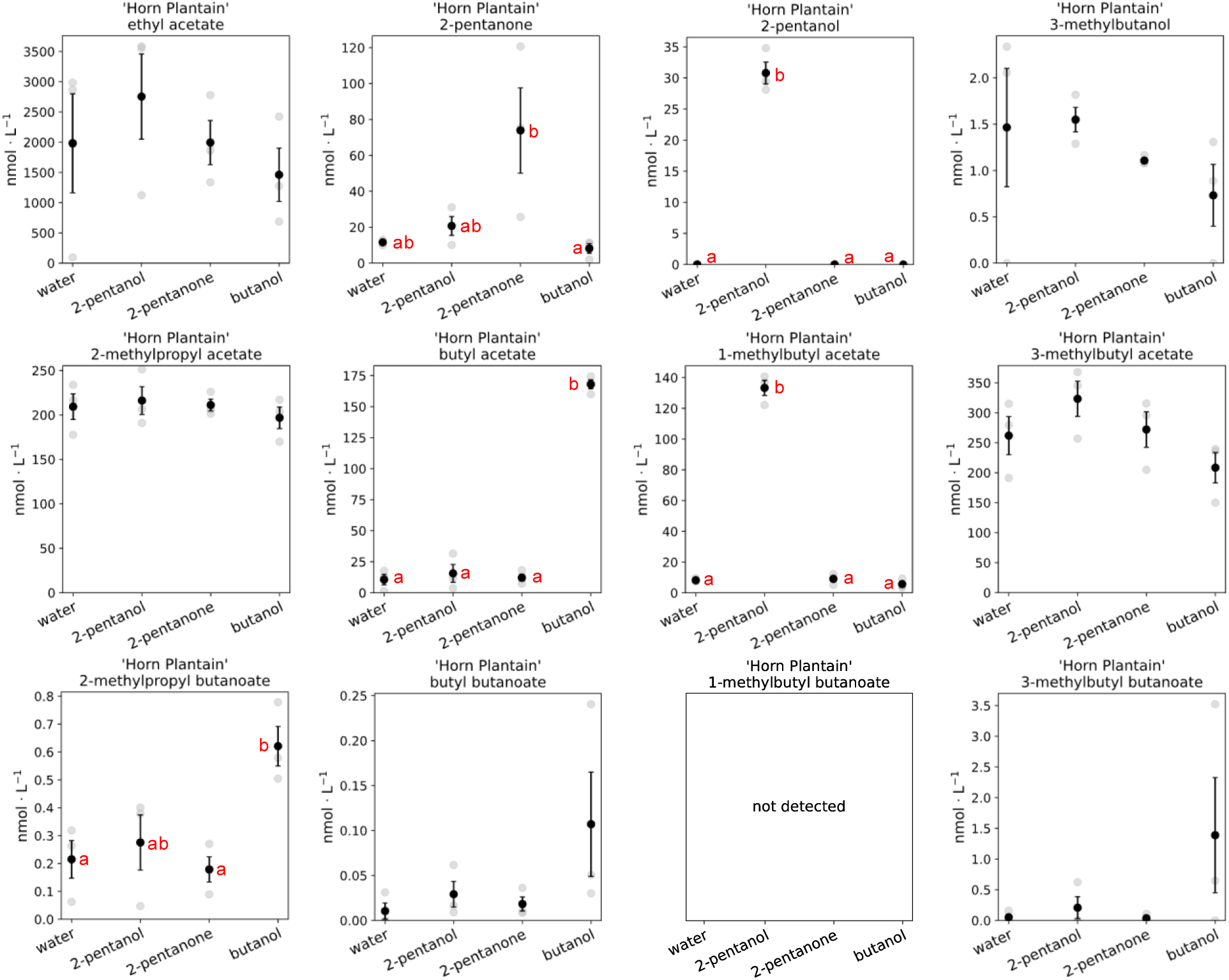
Volatile headspace concentrations of ‘Horn Plantain’ banana fruit pulp sections treated with potential aroma precursors. Presented as means ± ½ sᴅ of three biological reps. Significantly different volatile concentrations are denoted by different letters adjacent to means (Tukey’s test, ⍺=0.05).

**Figure S8.**
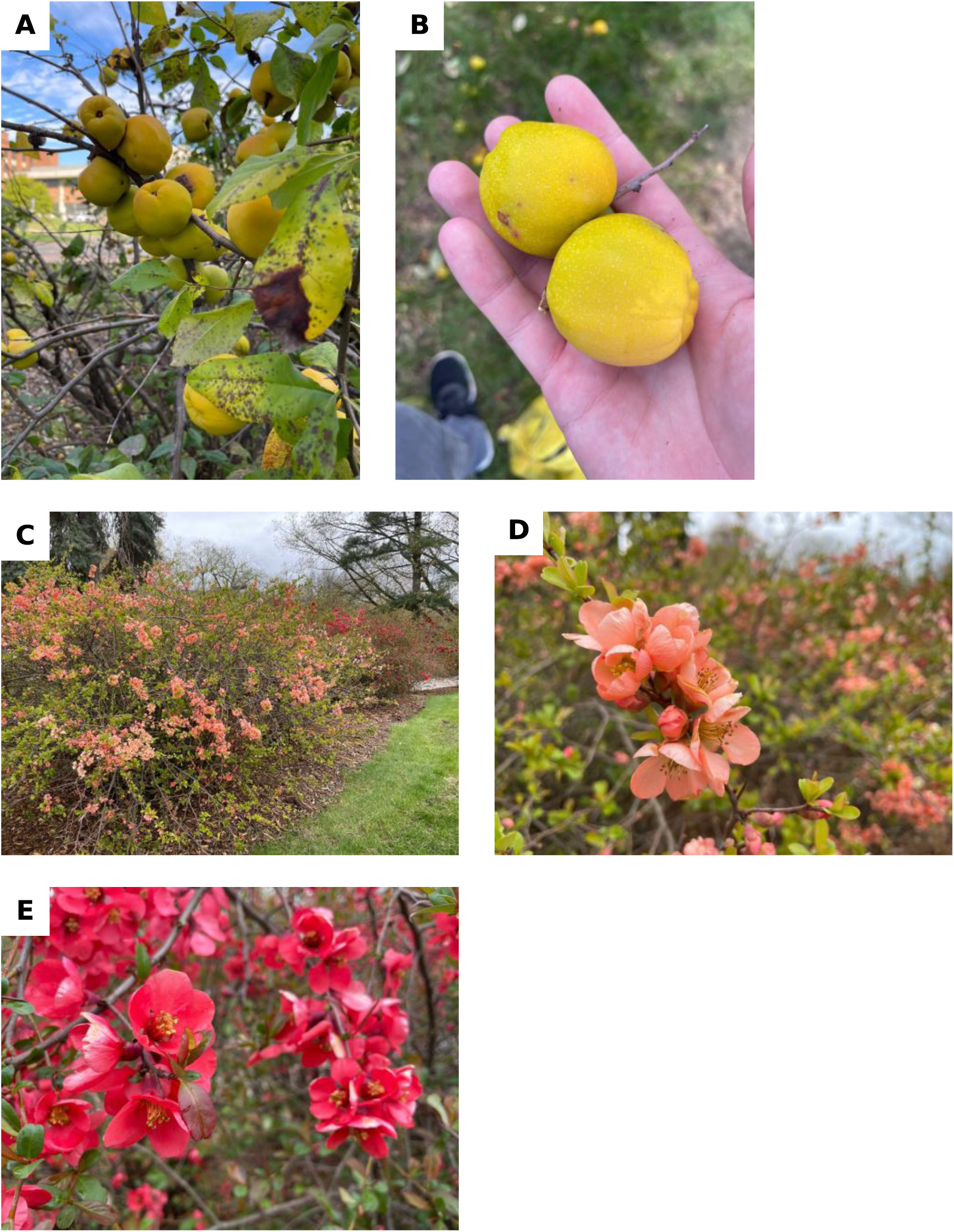
Photos of flowering quince (*Chaenomeles* ×*superba*). A-B) ‘Dr. Banks Pink’ fruit on October 6^th^, 2022. C-E) Flowers of flowering quince on May 10^th^, 2023. Pinkish flowers = ‘Dr. Banks Pink’; red flowers = ‘Crimson Beauty’ (not used in this study).

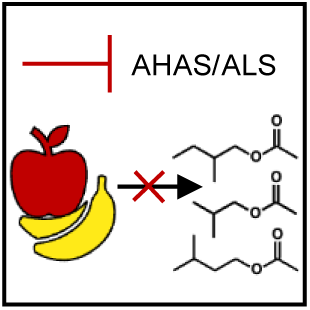
Table of Contents (TOC) Icon

## Supplemental Methods & Text

### Materials and methods

#### Plant material

‘Gala’, ‘Empire’, and ‘Jonagold’ apple (*Malus* ×*domestica* Borkh.) fruit were harvested from local orchards at commercial maturity and transported to the laboratory during the 2022 season. Developmentally, the fruit were at the onset of ripening but no aroma volatiles could be discerned subjectively. ‘Gala’ and ‘Empire’ fruits began treatment (see below) immediately after arrival. ‘Jonagold’ fruit were held in air at 0 ℃ for two days before transfer to 1.5 O_2_, 3% CO_2_, 0 ℃. ‘Jonagold’ fruit were held in these conditions for twelve days before the initiation of treatment.

Banana (*Musa* spp. AAA group, Cavendish subgroup, cv. Valery; *Musa* spp. AAB group, Plantain subgroup, cv. Horn Plantain) fruit that had not been treated with ethylene were obtained from a local supermarket produce distribution and ripening center (Meijer/Chiquita, Lansing, MI). ‘Horn Plantain’ fruit were held at room temperature (22 ℃) for 2-3 days before treatment. ‘Valery’ fruit were held at 13.5 ℃ until treatment, however fresher fruits provided more consistent results compared to older ones.

‘Dr. Banks Pink’ flowering quince (*Chaenomeles* ×*superba*) fruit were collected from accession CC7985*05 on the Michigan State University campus grounds. Fruits were actively producing aroma.

#### Treatment

Whole apple fruit were stored at room temperature (22 ℃) and rubbed daily with 3 mL of freshly made herbicide or water solution (1 mM rimsulfuron, made from DuPont™ Matrix® SG (25% w/w active ingredient), 0.1% Tween 20) before preparation with further treatments. Quince fruit had 2 mL applied daily.

‘Gala’, ‘Empire’, and ‘Jonagold’ apple fruit were treated for four, nine, or seven days before further preparation, respectively. Quince were treated for six days before analysis. Whole fruit routinely had their headspace volatiles sampled for aroma production and treatment efficacy to determine appropriate times for further treatments.

Acetate and ⍺-ketoacid feedings were performed by preparing vials of peel tissue as previously described by (Sugimoto, et al., 2021). In detail, 11 mm disks of peel from treated fruits were cut away from the fruit and trimmed to have 1-2 mm of cortex tissue using a cork borer and scalpel. Five peel disks were laid upon five filter paper disks of equal diameter resting upon a glass microscope slide trimmed to fit within a 22 mL glass vial. Each disk of filter paper was wetted with 20 µL of treatment solution, described below, before delicately placing a fruit tissue disk, peel side-up, upon the paper. More treatment solution, 20 µL, was then added on top of each peel disk before sliding the apparatus into the glass vial and sealing with a Mininert valve (Thermo Scientific). Samples were incubated overnight (22 ℃) before headspace sampling.

‘Gala’ fruit were fed with 10 mM methanol, 10 mM acetate (unlabeled or 1,2-^13^C_2_) pH 7 balanced with KOH, 0.05% Tween 20, and 0.5 mM rimsulfuron if appropriate. These concentrations were doubled for ‘Empire’ and ‘Jonagold’ fruit and contained, when appropriate, 20 mM of ⍺-keto-β-methylvalerate, ⍺-ketoisovalerate, or ⍺-ketoisocaproate.

Banana fruit were prepared via two means, both inspired by past studies (Palmer & McGlasson, 1969). The first method was used for studying the effects of sulfonylureas on volatile and amino acid content, and for the feeding study with ‘Horn Plantain’ fruit, whereas the second was used to investigate the recovery of sulfonylurea-treated fruits after ⍺-ketoacid application.

In the first method, mature green ‘Valery’ fruit were cut into ∼1 cm^2^ squares from fruit not treated with ethylene. Care was taken such that one edge of the square was from the edge of the pulp, thus representing an undisturbed or ‘live edge’ of cells that should be able to maintain unhindered gas exchange. Five of these pulp sections were placed on filter paper disks on a trimmed microscope slide and placed into 22 mL vials as above described for apples. Each pulp section and piece of paper also received 20 µL of treatment solution with or without herbicide [0.5 mM methyl-halosulfuron (referred herein as halosulfuron, made from Sandea® (75% w/w active ingredient), 0.1% Tween 20]. The vials were sealed with Mininert valves. Fruit ripening was then induced by injecting 1 µL ethylene into the vials. The following day the vials were vented for 15 min before having the Mininert replaced on the vial but with a needle inserted to allow for gas diffusion. The following two days, 20 µL of freshly made herbicide or water solution was added onto the pulp. The needles were kept in the Miniert valves to maintain gas diffusion. The next day, and thus four days since initially preparing the vials, the needles were removed and the vials were sealed and incubated for at least 1 hour at room temperature (22 ℃) before headspace sampling.

Plantains were prepared in a similar way but did not receive herbicide. Before analysis (∼1-2 hours), 10 µL of water, 1 mM butanol, 1 mM 2-pentanone, or 1 mM 2-pentanol were added to the pulp sections.

While the results of inhibition using this methodology were consistent in ripening fruits, maintaining the small pulp sections at an appropriate humidity to allow for ripening while also preventing desiccation and microbial outbreak proved to be exceedingly difficult. Thus a second methodology that more closely followed the work of (Palmer & McGlasson, 1969) and utilized much larger transverse cross-sections of the fruit was used for further experimentation. This second method, when fresh fruit were used, led to more reliable ripening with less maintenance than the first method as well as the same volatile content shifts.

In detail, mature green ‘Valery’ fruit that had not been treated with ethylene were first surface sterilized with a spray of 75% v/v ethanol before continuing preparation in a sterile laminar flow hood. Transverse cross-sections of the intact fruit, including peel and pulp, were prepared with an industrial onion cutter set to 5 mm thickness. The slices were labeled on the peel with permanent marker before being submerged for ∼5 min in water. The slices were then submerged in 0.1% Tween 20, with or without 0.5 mM halosulfuron, in a 200 mL glass Mason jar, with ∼50 mL of air in the jar’s headspace. The slices were then vacuum infiltrated for 2 min before being patted dry. Afterwards, the slices were then placed on a scaffold made from egg crate light diffuser panels within a 10 L desiccator with ∼200 mL of water in the base. The slices were then incubated with 125 µL · L^-1^ ethylene within the closed desiccator. After incubation, 0.5 L · min^-1^ of hydrated air was then on supplied to the desiccator. Five days after slicing the fruits, 0.5 mL of treatment solution was applied to the surface of the slices. Solutions consisted of 0.1% Tween 20 and, in various combination, 0.5 mM halosulfuron, 20 mM ⍺-keto-β- methylvalerate, 20 mM ⍺-ketoisovalerate, and 20 mM ⍺-ketoisocaproate. The resulting aroma profiles were sampled on the following day.

#### Volatile analysis

Headspace volatiles from vials were sorbed for 30 s using a solid-phase micro extraction (SPME) fiber (65 μm PDMS-DVB; Supelco Analytical, Bellefonte, PA). The SPME fiber was then directly desorbed for 1 min in the injection port of a gas chromatograph (GC; HP-6890, Hewlett-Packard, Wilmington, DE) coupled to a time of flight mass spectrometer (MS; Pegasus II, LECO, St. Joseph, MI). Desorbed volatiles were cryofocused at the beginning of the column by immersing said region of the column in liquid nitrogen. After the desorption period, the run was initiated and the liquid nitrogen removed.

Quince fruits selected for volatile analysis and banana slices were incubated for 20 min at room temperature (22 ℃) in 1 L sealed Teflon jars before a 3 min sorption and 2 min desorption protocol as described above.

The conditions of the system were as follows. Injection port: 200 ℃, splitless, helium carrier gas, front inlet flow was 1.5 mL · min^-1^ constant, 10 mL · min^-1^ purge flow, 11.5 mL · min^-1^ total flow. Oven: initial temperature at 40 ℃ for 0 min, ramped by 43 ℃ · min^-1^ to 185 ℃ for 0 min. Column: HP-5MS, 30 m × 0.25 mm i.d., 0.25 µm film thickness (Agilent, Santa Clara, CA). Transfer line temperature was 225 ℃. MS: Electron ionization (-70 eV), ion source temperature was 200 ℃, solvent delay was 50 sec, m/z 29 to 400 were scanned for, detector voltage was 1500 V, data collection rate was 20 Hz.

When ⍺-ketoisocaproate was supplied to the fruit and separation of 2-methylbutyl acetate and 3-methylbutyl acetate was of interest, the following oven parameters were used: initial 40 ℃ for 0 min, 10 ℃/min until 100 ℃, 20 ℃ · min^-1^ until 130 ℃, 60 ℃/min until 185 ℃.

Compounds were identified by comparison with the retention time and mass spectrum against authenticated reference standards and spectra (National Institute of Standards and Technology Mass Spectral Search Program Version 2.0, 2001). Volatiles were quantified by calibration with a standard of 59 authenticated compounds (Sigma-Aldrich Co., St. Louis, MO and Fluka Chemika, Seelza, Germany). The standard was made by placing 0.5 μL of an equal- part mixture of the neat compounds onto a disc of filter paper before quickly placing the filter paper into a 4 L sealed flask fitted with a Mininert valve (Valco Instruments Co. Inc., Houston, TX) for SPME fiber access. The quantification m/z of each compound can be seen in Supplemental Table S13.

After volatile analysis, apple peel disks, collected peel of quince, and ‘Valery’ pulp section samples were held at -80 ℃ for further amino acid analysis.

#### Sensory analysis

‘Jonagold’ fruit, previously stored in CA, as described above, were treated for seventeen consecutive days with herbicide or water solution (1 mM rimsulfuron, made from Matrix®, 0.1% Tween 20) as described above. Fruits were screened via GCMS, described above, to ensure successful treatment and suppression of 2-methylbutyl and 2-methylbutanoate esters. Treatment was stopped two days in advance of our initial target date for the sensory analysis and the fruit were rinsed with warm water to remove residual herbicide solution. However, the experiment was delayed due to the February 13, 2023, mass shooting at Michigan State University. The fruit, having been treated and screened, were held in air at 1 ℃ under plastic bags seven days as the campus community recovered from the tragedy. The day before the new sensory study date, the fruit were removed from storage and allowed to warm to room temperature (22 ℃). The morning of the experiment, 1 x 5 cm segments weighing 2.5 – 3 g, were prepared from the apples. Cortex tissue was trimmed to maximize the proportion of peel tissue. Segments of fruit were prepared instead of discs to limit the surface area of cut tissues, minimizing oxidative volatiles. Fruit segments were then placed in 40 mL amber vials and sealed with PTFE-lined caps. Samples were allowed to incubate at least 2 hours before sensory evaluation. All evaluations were conducted within a 3-hour period to minimize changes in sample aroma. Each vial was opened no more than once per hour, and vials were used a maximum of three times during the study period.

Study participants were recruited from the campus of Michigan State University and the greater Lansing, MI area. Participant demographics were as follows: N = 20; age: (range = 18 – 41, average = 24.4); gender: (14 female, 6 male); sex assigned at birth: (14 female, 6 male); race:(1 African American or Black, 7 Asian (3 Asian Indian, 3 Chinese, 1 Vietnamese), 12 Caucasian or White); ethnicity: (17 Not of Hispanic, Latino/a or Spanish origin, 1 Mexican American, 1 unknown, 1 Spanish). Eligibility criteria were as follows: 1) be willing to smell fresh fruit, 2) have a normal sense of taste and smell, 3) be between the ages of 18-55. Subjects were compensated with a $10 gift card voucher for their participation. This protocol was reviewed and determined to qualify for exempt status by the Michigan State University Institutional Review Board (Study 00008470).

Participants performed twelve Duo-Trio trials, each with a blinded treated and untreated sample, labeled with randomized 3-digit codes, as well as an untreated sample labeled as the reference. The presentation order of the blinded samples followed a randomized complete block design. Participants were prompted: “In front of you is a set of three samples. Smell the reference sample labeled REF and then smell the two test samples. Select the sample code that smells the same as the REFERENCE sample. You must make a choice, even if it is only a guess. You may re-smell as often as you wish.” In between each trial, there was an enforced 60-second break. The sensory survey was built and deployed using RedJade software (RedJade Sensory Solutions, LLC). Testing took place in sensory booths to provide a controlled environment and minimze biasing interactions. Binomial statistics were performed with Microsoft Excel v16.69.1. p-value = 1-BINOM.DIST(correct responses-1,total trials, 0.5, TRUE).

#### Citramalate synthase sequencing

Leaves of accessions of the USDA Geneva *Malus* Core Collection that have previously been identified as being first-degree relatives of ‘Cox’s Orange Pippin’ were kindly collected, frozen, and shipped to our laboratory on dry ice by the staff of the USDA, ARS, Plant Genetic Resources Unit at Geneva, NY. Samples were held at -80 ℃ prior to being ground to a powder in in liquid nitrogen-chilled mortar and pestles. DNA was extracted with DNeasy Plant Mini Kit (Qiagen, Hilden, Germany). The surrounding locus of the G → C SNP that results in a change of the 387^th^ codon of GAG → CAG, and thus a change of the amino acids Gln → Glu was amplified via PCR. The conditions of the 20 µL were as follows: 1X GoTaq® Green Master Mix (Promega, Madison, WI), 0.25 µM forward primer (GTGGAAGAGTACAGCGGATT), 0.25 µM reverse primer (GCCAAAATAATCTCATAGGTGCTC), ∼200 ng DNA, balanced with water. The reaction conditions were as follows: 2 min 95 ℃, followed by 30 cycles of 30 s 95 ℃, 30 s 51 ℃, 20 s 72 ℃, followed by 5 min 72 ℃. Products were verified with gel electrophoresis prior to purification with Monarch® PCR & DNA Cleanup Kit (NEB, Ipswich, MA). The amplicon was Sanger sequenced with both primers at the Research Technology Support Facility Genomics Core at Michigan State University.

#### Amino acid analysis

Frozen samples were ground to a powder in liquid nitrogen-chilled mortar and pestles. About 0.5 mg of tissue were vortexed for 10 sec in 2 mL of room temperature (22 ℃) 1:1:1 (water:acetonitrile:ethanol, v/v) spiked with 2 nmoles of U-^13^C,^15^N labeled amino acids (MilliporeSigma) before being heated for 15 min in a 65 ℃ water bath. Extracts were then briefly chilled on ice before being centrifuged at 4400 × g for 15 min at 4 ℃. The supernatant was filtered by centrifugation (0.2 μm nylon centrifugal filter; Costar, Corning) at 21000 × g for 5 min at room temperature. 10 µL of the filtrate was transferred to an autosampler vial and diluted 100-fold with 990 µL of 10.1 mM perfluorohexanoic acid (PFHA) spiked with 2 µmoles of internal standard. Thus the final concentration of internal standard was ∼2 µM.

An amino acid standard series was prepared from a premade mixture (Millipore Sigma, AAS18) that contained equal molar amounts of cystine and all 20 proteinogenic amino acids save for tryptophan, asparagine, glutamine, and cysteine. An equal molar mixture of tryptophan, asparagine, glutamine, and cysteine was subsequently prepared. To avoid dilution errors or artefacts from differing buffers, these amino acid stocks were aliquoted and desiccated such that a five-part standard series ranging from 250 µM to 25 nM would be produced upon resuscitation with 10 µL of spiked extraction buffer and 990 µL spiked PFHA solution.

Samples and amino acids were held overnight at -20 ℃ before analysis.

Amino acids were analyzed with a Xevo TQ-S Micro UPLC (H-Class)-MS/MS (Waters, Milford, MA) at the Michigan State University Mass Spectrometry and Metabolomics Core. Conditions were as follows. HPLC column: Acquirt UPLC HSS T3, 2.1 x 100 mm, 1.7 µm particle size (Waters), with a 0.2 µm pre-column filter (Waters). Mobile phase: A) 10 mM PFHA in water, B) acetonitrile. LC gradient: linear gradient, slope setting = 6, flow rate = 0.3 mLᐧ min^-1^, step 1) 0 min, 100% A, 0% B, 2) 1 min, 100% A, 0% B, 3) 8 min, 35% A, 65% B, 4) 8.01 min, 10% A, 90% B, 5) 9 min, 10% A, 90% B, 6) 9.01 min, 100% A, 0% B, 7) 13 min, 100% A, 0% B. Column temp: 40 ℃. Autosampler temp: 10 ℃. Injection volume: 10 µL. Tune parameters: electrospray ionization, standard ESI probe, capillary voltage = +1.0 kV, source temp = 120 ℃, desolvation temp = 350 ℃, desolvation gas = 800 Lᐧ hr^-1^, cone gas = 40 Lᐧ hr^-1^. MS collection was split into three phases and were adjusted after checking the retention time of several samples, however proline was missed for apple and quince samples. Parent and daughter ions, cone and collision voltages, phases collected and approximate retention times can be seen in Supplemental Table S14.

Data were quantified by first calculating a linear regression of log(unlabeled amino acid response/labeled amino acid response) transformed standard responses. R^2^ values were all greater than 0.98 and the slope (m) and y-intercept (b) were used to calculate unknowns: µM of unknown sample = 10*[(log(unknown unlabeled response/unknown labeled response)-b/m].

#### Population genetics of citramalate synthase

The ability of panelists to discriminate the lack of 2-methylbutyl and 2-methylbutanoate esters in a complex aroma profile led us to consider what implications this may have had on apple breeding and the prevalence of these esters among commercially grown cultivars.

It has been demonstrated that the presence of an active allele of citramalate synthase is necessary for apples to synthesize a copious amount of 2-methylbutyl and 2-methylbutanoate esters in a dominant/recessive phenotypic relationship (Sugimoto, et al., 2021) (Sugimoto, et al., 2015). Among the 99 cultivars from the USDA Geneva *Malus* Core Collection previously analyzed, 6.1% were homozygous recessive, 36.4% were heterozygous, and 57.6% were homozygous dominant. Apples that were homozygous recessive had a significantly lower ratio of 2-methylbutyl and 2-methylbutanoate esters to straight-chain esters as compared to those with at least one copy of the active allele. From this data it cannot be determined if the observed allelic distribution favoring branched-chain ester-producing phenotypes is a result of natural or artificial selection.

However, among the apples screened, ‘Cox’s Orange Pippin’, which has been identified through pedigree and sequencing-based analyses as a common breeding parent to many cultivars, was found to be homozygous recessive for citramalate synthase (Sugimoto, et al., 2021) (Muranty, et al., 2020) (Noiton & Alspach, 1996). Through sequencing-based pedigree analyses of the USDA Geneva *Malus* Core Collection, several dozen cultivars with a first-degree (parent-offspring or clonal) relationship with ‘Cox’s Orange Pippin’ have been identified (Migicovsky, et al., 2021). If no selective pressure were occurring, then, given that ‘Cox’s Orange Pippin’ is homozygous recessive, a relatively large proportion of its offspring will likewise be homozygous recessive and unable to produce 2-methylbutyl and 2-methylbutanoate esters.

More than half of the cultivars within the USDA Geneva *Malus* Core Collection have been found to be interconnected through first-degree relationships, suggesting a large degree of interbreeding and the reasonable assumption that the aforementioned distribution of alleles observed by Sugimoto et al. (2021) is representative of the population of ‘Cox’s Orange Pippin’s potential mates. We reasoned that if the proportion of ‘Cox’s Orange Pippin’ offspring that are homozygous recessive be less than expected, assuming a parent population akin to previous observation, then it is likely that humans have, likely unknowingly, been selecting for cultivars that are able to synthesize copious amounts of anteiso-branched-chain esters as opposed to cultivars that lack this ability.

We sequenced the consequential single nucleotide polymorphism of citramalate synthase from 40 cultivars previously identified as having a first-degree relationship with ‘Cox’s Orange Pippin’ and termed this as the offspring population (Migicovsky, et al., 2021). The parent population (from Sugimoto et al. (2021)) and offspring population were mutually exclusive save for ‘James Grieves’, however this cultivar has been identified as being an offspring of ‘Cox’s Orange Pippin’ and was thusly sequestered to the offspring population (Muranty, et al., 2020). Two cultivars, ‘Cherry Cox’ and ‘Potter Cox’, were identified as being sport mutations of ‘Cox’s Orange Pippin’ whereas another two cultivars, ‘Margil’ and ‘Rosemary Russet’, have been determined to be the parents of ‘Cox’s Orange Pippin’ (Muranty, et al., 2020) (Howard, et al., 2023). The sports (both homozygous recessive), and parents (both heterozygous) were removed from further analysis. In keeping with population definitions set by Migicovsky et al. (2021), cultivars listed as recently derived hybrids were removed from the parent and offspring populations as well. Furthermore, cultivars listed by alphanumeric codes were also removed. It was reasoned that these cultivars, given their lack of a common name and their status only as breeding lines, have not been publicly released and are thus not likely to be considered to be of high enough quality for commercial success.

Given our parent population, when crossed with ‘Cox’s Orange Pippin’ we would expect 29.1% of offspring to be homozygous recessive if no selection is occurring. However, among the 32 cultivars of the ‘Cox’s Orange Pippin’ offspring population, only 3 (9.4%) were homozygous recessive (p = 0.0141).

